## Supplement 1 for "*In vivo* chromatic and spatial tuning of foveolar retinal ganglion cells in *Macaca fascicularis*"

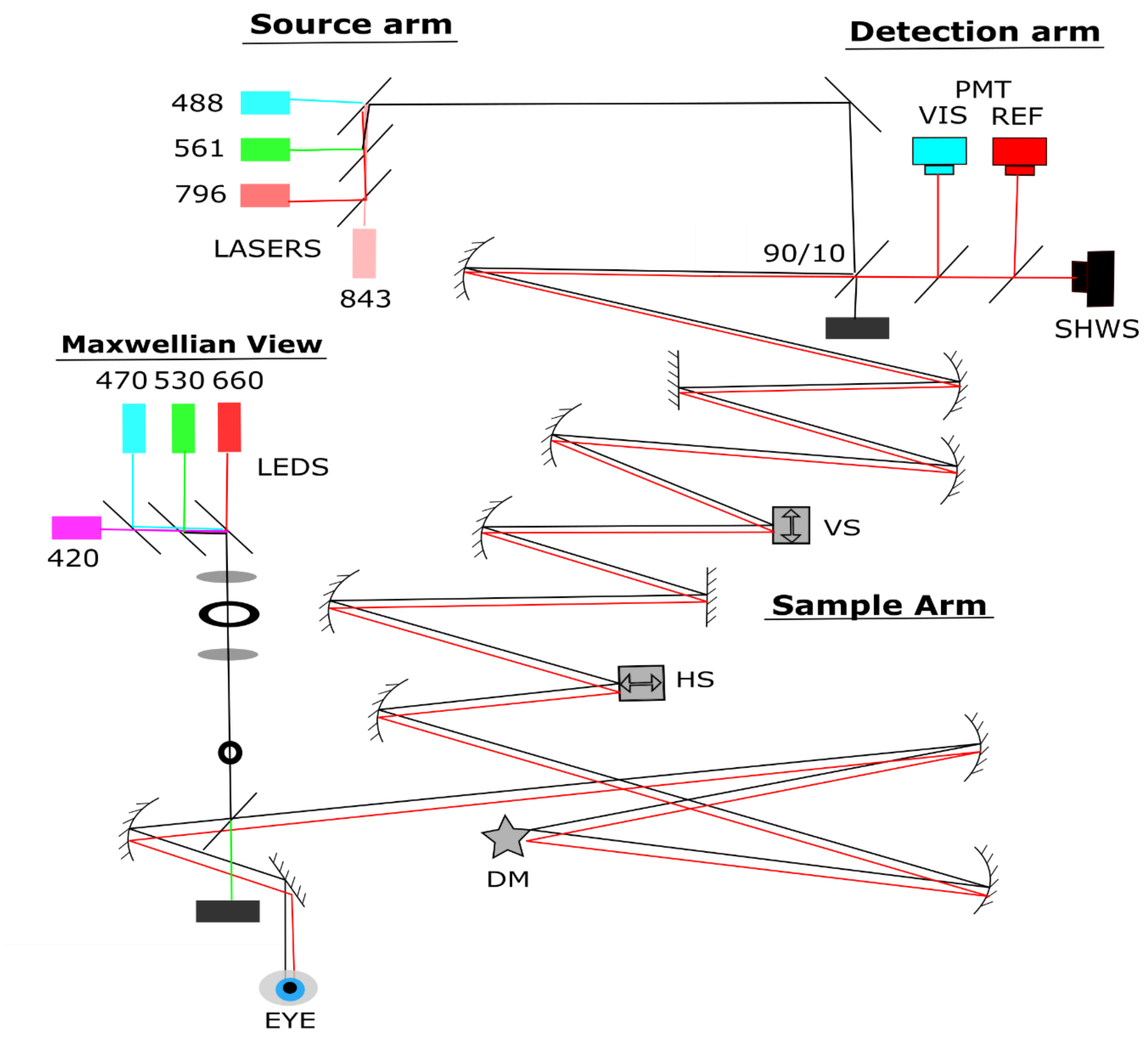


**S1 Fig. AOSLO System Diagram.** The adaptive optics scanning light ophthalmoscope used in this study consists of three main arms or subsystems. First, the Source arm, contains the 488 nm laser used to excite GCaMP fluorescence, the 561 nm laser used as visual stimulation as in the drifting gratings, the 796 superluminescent diode used to image the cone photoreceptors, and the 843 nm laser diode used for wavefront sensing and correction. Each source is co-aligned and combined with dichroic mirrors so that at the output of the arm they exit coaxial with each other along the same beam path into the system. 90% of the source power is lost at a 90/10 beamsplitter such that 10% of the source power enters the Sample arm of the system—this is done so that on the back-pass, 90% of the signal is transmitted to the detectors while only 10% of the signal is lost. The Sample arm is composed of various optical telescopes (whose function is to translate the beam axially and change the magnification to match the pupil sizes of the various active elements) and four important planes conjugate to the pupil. The first two conjugate planes are the vertical (VS) and horizontal (HS) scanners which raster scan the beam across a rectangular field of view. The third conjugate plane is the deformable mirror (DM) which changes shape to correct the measured wavefront aberrations of the animal's eye to provide near-diffraction limited imaging of the retina. The fourth conjugate plane is the pupil of the animal's eye, where all source light enters and from which all detected light emanates. In the Detection arm, two photomultiplier tubes (PMT) detect visible fluorescence (VIS) from the emitted GCaMP signal and infrared (REF) reflected light from the cone photoreceptors. A Shack-Hartmann wavefront sensor (SHWS) collectes infrared light from the 843 nm source and measures the wavefront aberrations of the animal's eye, sending that information to the deformable mirror in a closed loop. Finally, a Maxwellian View subsystem is used to inject the LED stimulation lights into the system near the pupil plane (bypassing both scanners and the deformable mirror). The subsystem contains LEDs at 420 nm, 470 nm, 530 nm, and 660 nm, although the 470 nm LED was not used in this study. Just as in the Source arm, the LEDs are co-aligned and combined using dichroic mirrors so they follow the same beam path. The LEDs pass through a spatial filter which removes any spatial inhomogeneities and then through a field stop which restricts the beam sizes on the retina to a circular subsection of the fovea about 1.3 degrees in diameter. The LEDs are reflected into the system via a pellicle beamsplitter which has 88% transmission (throws away 88% of the LED source power, but allows 88% of the signal to pass on the way to the detectors).
