## Supplement 2 for "*In vivo* chromatic and spatial tuning of foveolar retinal ganglion cells in *Macaca fascicularis*"

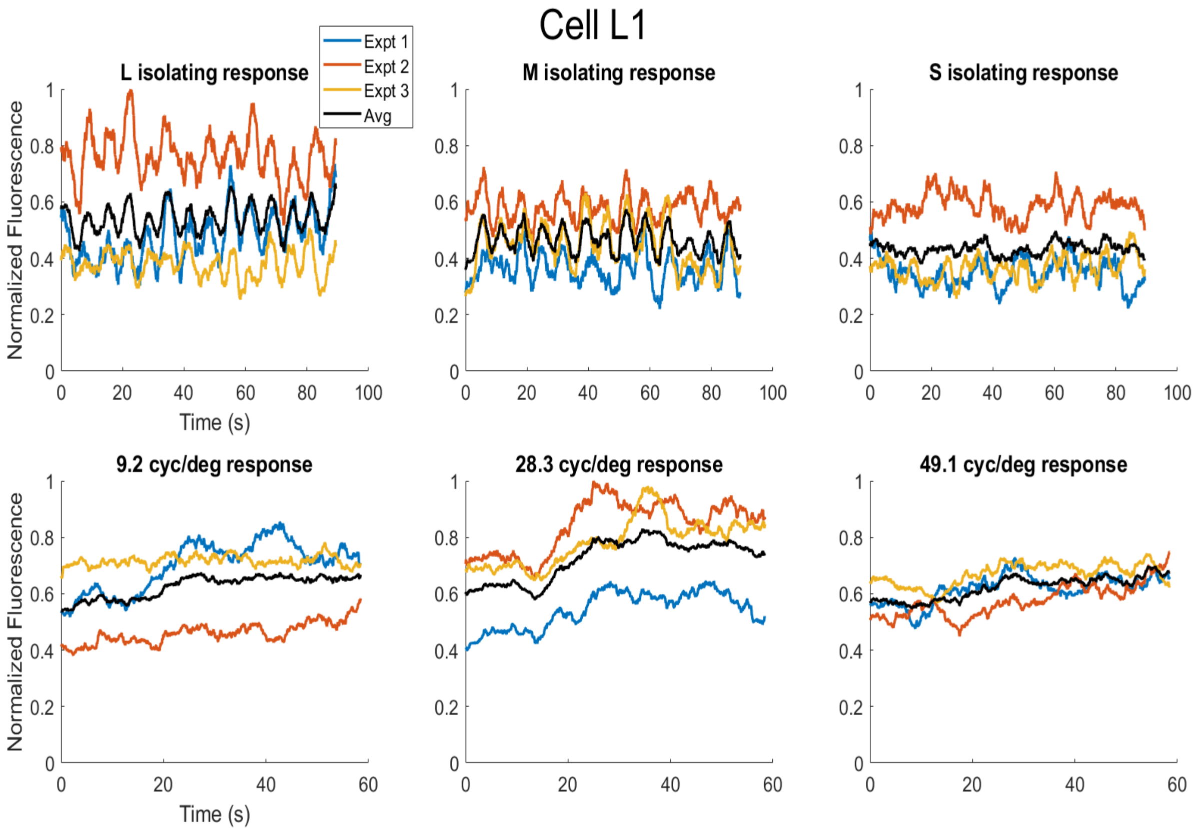
Cell L1
Expt 1
Expt 2
Expt 3
Avg
L isolating response
M isolating response
S isolating response
9.2 cyc/deg response
28.3 cyc/deg response
49.1 cyc/deg response
Normalized Fluorescence
Time (s)

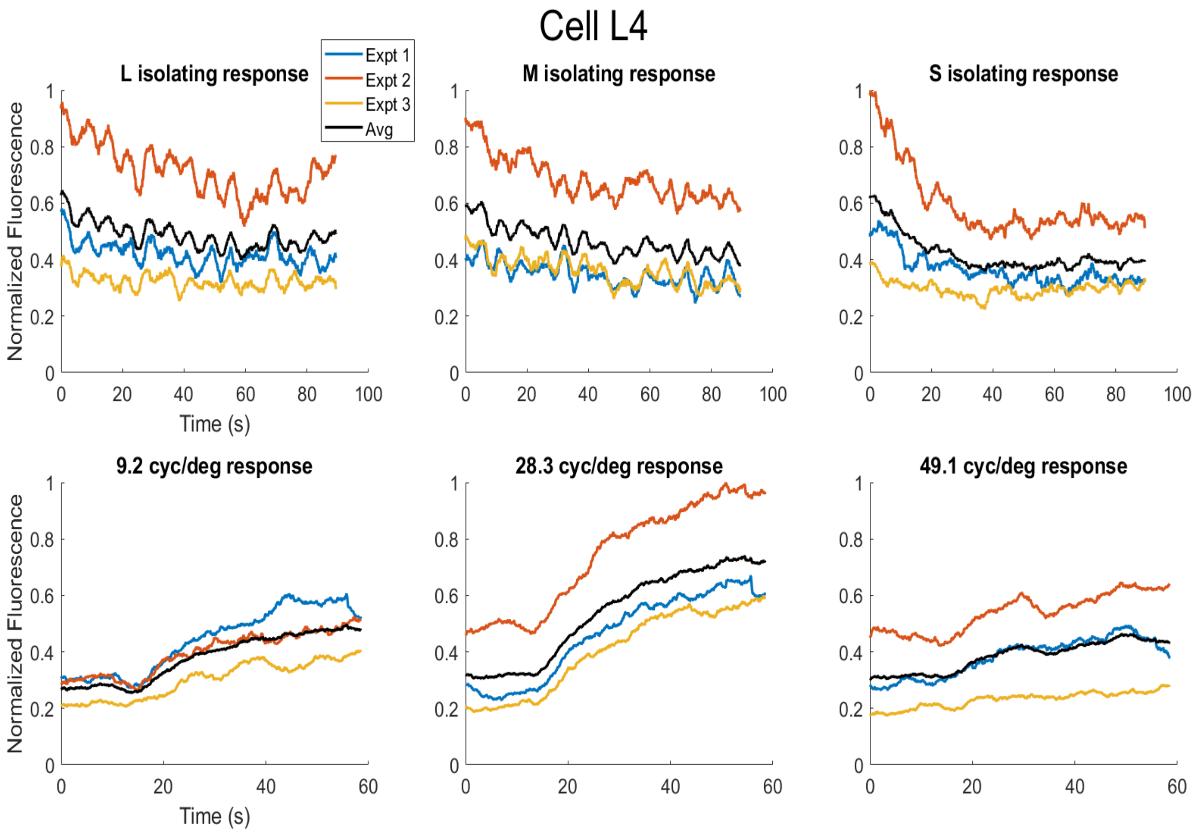
Cell L4
Expt 1
Expt 2
Expt 3
Avg
L isolating response
M isolating response
S isolating response
9.2 cyc/deg response
28.3 cyc/deg response
49.1 cyc/deg response
Normalized Fluorescence
Time (s)

### Cell L7

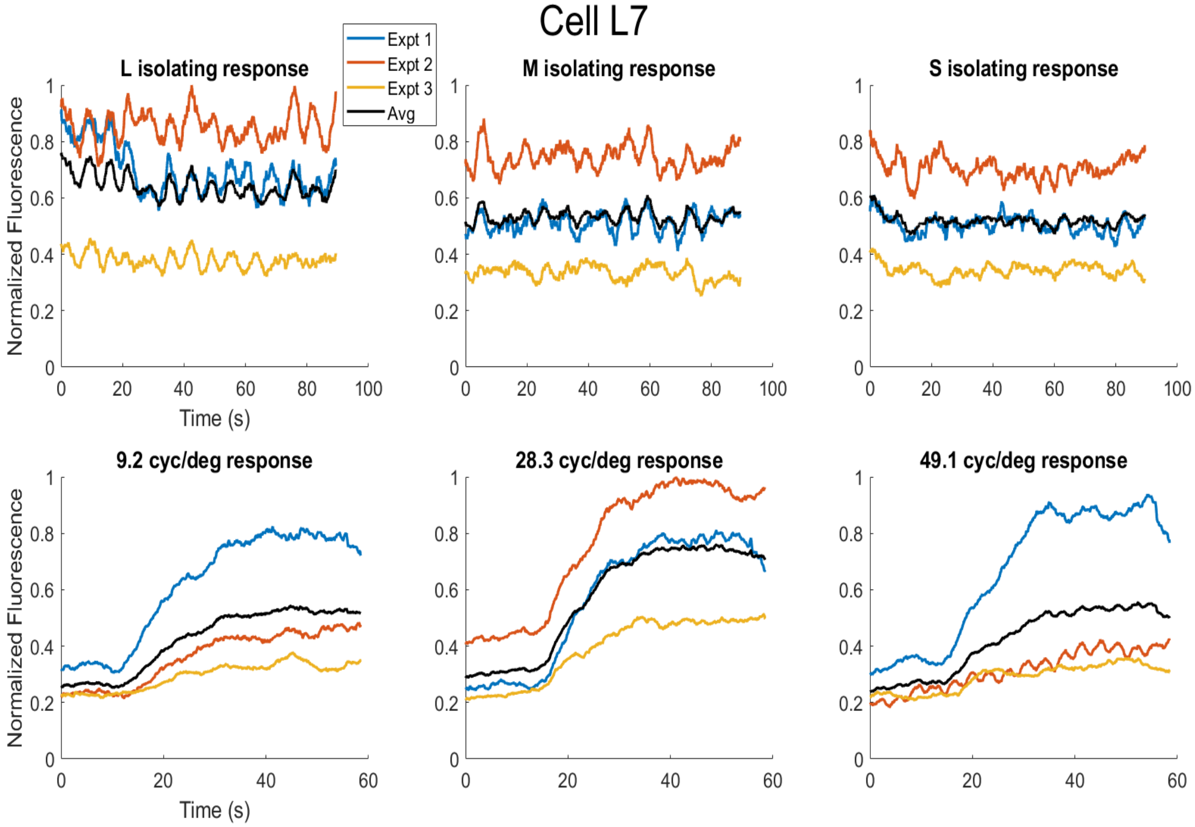


### Cell M3

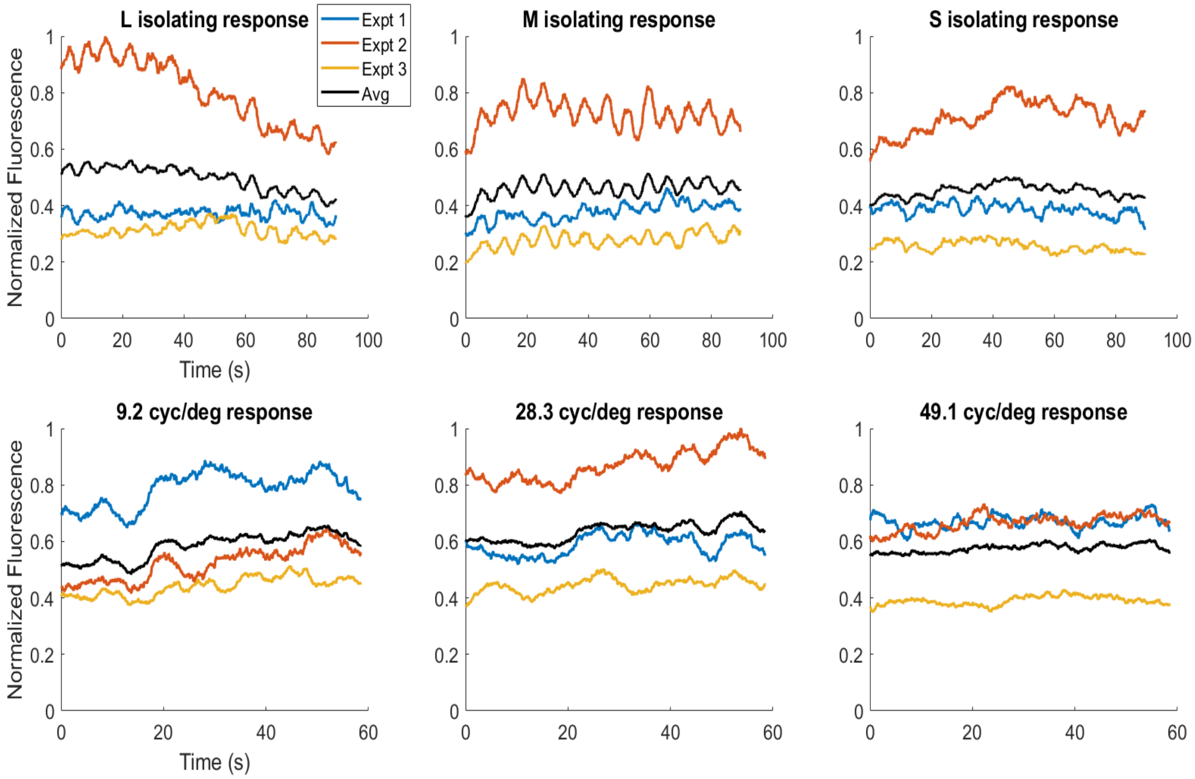

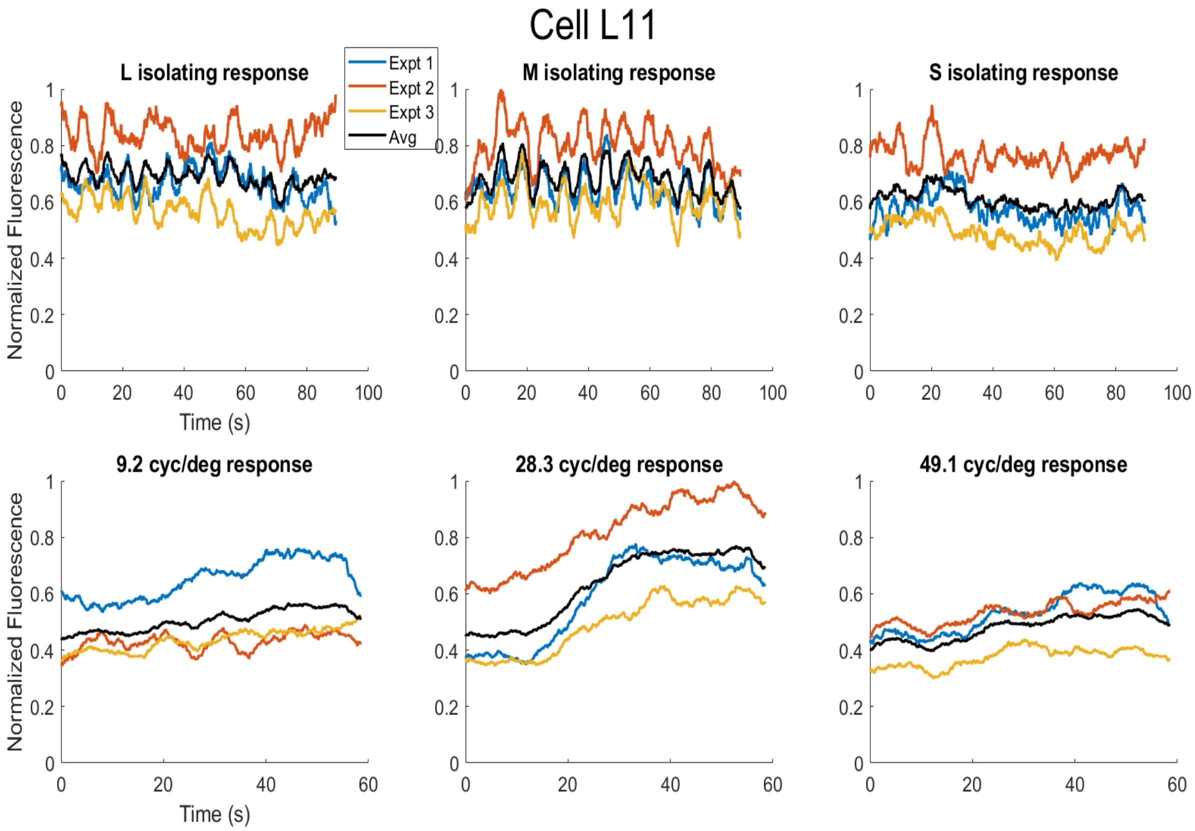
Cell L11
L isolating response
M isolating response
S isolating response
Expt 1
Expt 2
Expt 3
Avg
9.2 cyc/deg response
28.3 cyc/deg response
49.1 cyc/deg response
Normalized Fluorescence
Time (s)

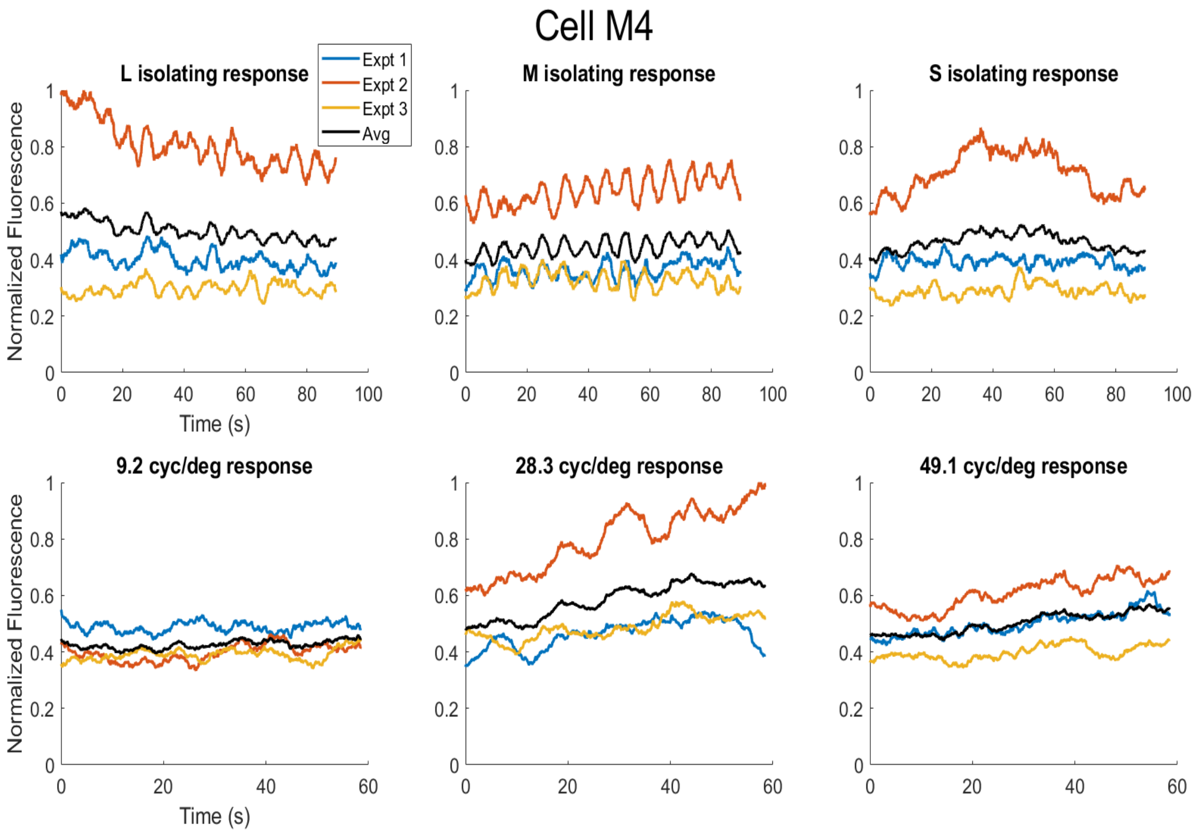
Cell M4
L isolating response
M isolating response
S isolating response
Expt 1
Expt 2
Expt 3
Avg
9.2 cyc/deg response
28.3 cyc/deg response
49.1 cyc/deg response
Normalized Fluorescence
Time (s)

**S2 Fig. Examples of raw fluorescence time courses of putative midget cells to various stimuli.** Six of the putative midget cells identified from the foveola of animal M3 are shown in detail. For each cell, the fluorescence time course is shown for that cell for six different stimuli: the L-isolating 0.15 Hz flicker, the M-isolating 0.15 Hz flicker, the S-isolating 0.15 Hz flicker, the 9.2 cyc/deg 6 Hz drifting grating, the 28.3 cyc/deg 6 Hz drifting grating, and the 49.1 cyc/deg 6 Hz drifting grating. For each stimulus, there are three fluorescence traces corresponding to experiment one (blue), experiment two (orange), and experiment three (yellow), which all occurred approximately a week apart. There is a fourth fluorescence trace (black) that is the average over the three experiments shown. Each fluorescence time course was normalized to the peak response for that cell, and a moving window (MATLAB movmean()) was used to smooth each fluorescence time course with a width of three seconds for the L/M/S isolating stimuli and a width of 5 seconds for the drifting grating stimuli. Note that as described in the manuscript, the L and M isolating stimuli are characterized by a modulation of the fluorescence near the stimulus frequency of 0.15 Hz. As these six cells were all putative midgets, note that the S isolating responses do not have the same modulation frequency, and are mainly affected by the respiration rate of the animal which causes small residual motion of the eye (approximately 0.28 Hz in M3). Note that as described in the manuscript, the drifting grating responses are characterized by a sustained increase in the mean fluorescence as opposed to a modulation, since the drifting speed exceeded the GCaMP6 temporal sensitivity.
