## Supplement 3 for "*In vivo* chromatic and spatial tuning of foveolar retinal ganglion cells in *Macaca fascicularis*"

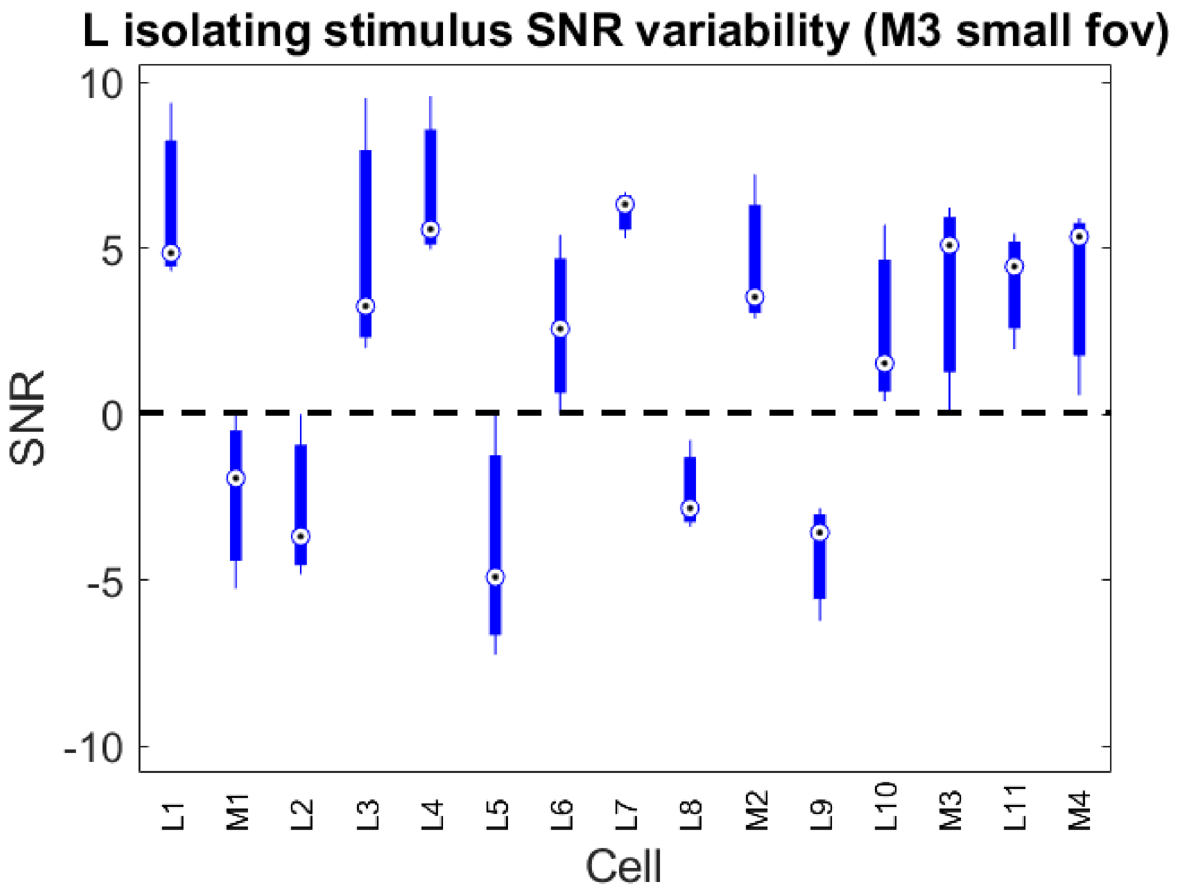
L isolating stimulus SNR variability (M3 small fov)
10
5
0
-5
-10
SNR
L1
M1
L2
L3
L4
L5
L6
L7
L8
M2
L9
L10
M3
L11
M4
Cell

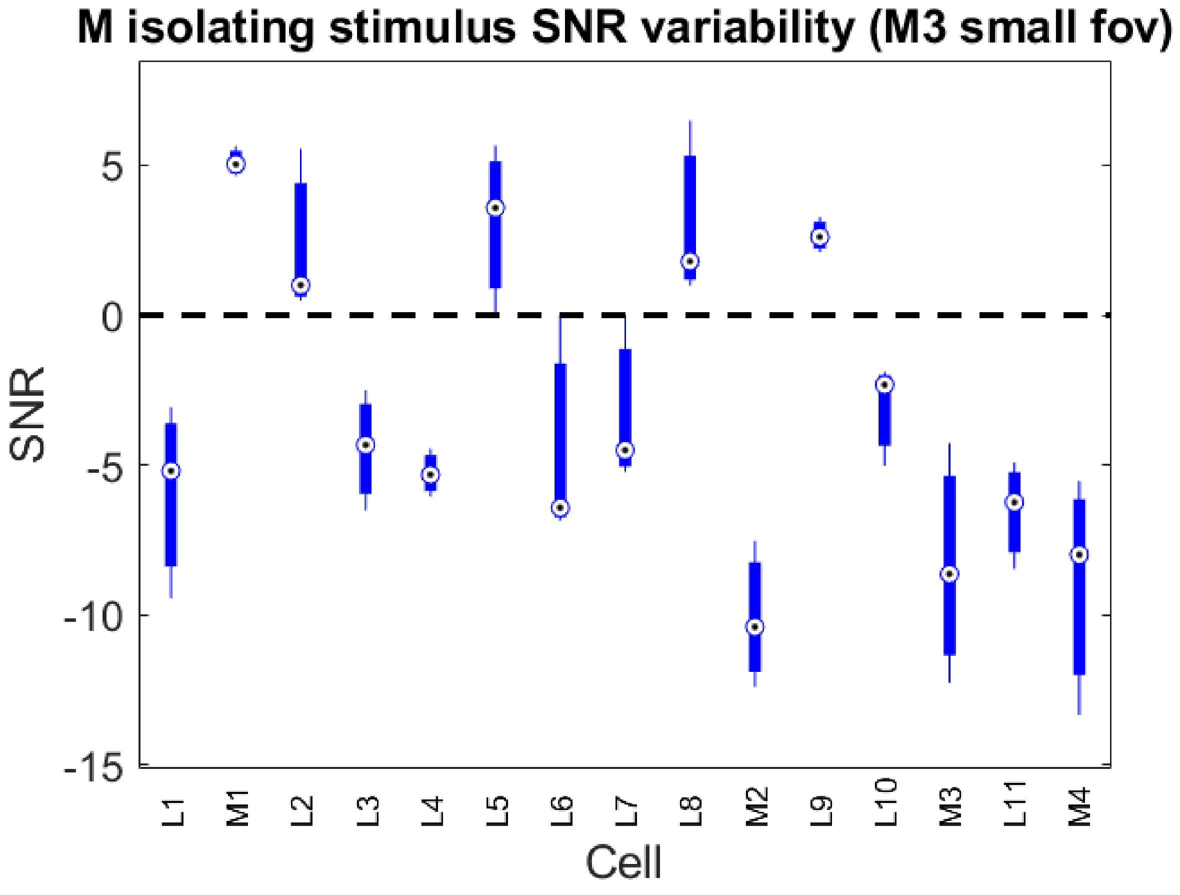
M isolating stimulus SNR variability (M3 small fov)
5
0
-5
-10
-15
SNR
L1
M1
L2
L3
L4
L5
L6
L7
L8
M2
L9
L10
M3
L11
M4
Cell

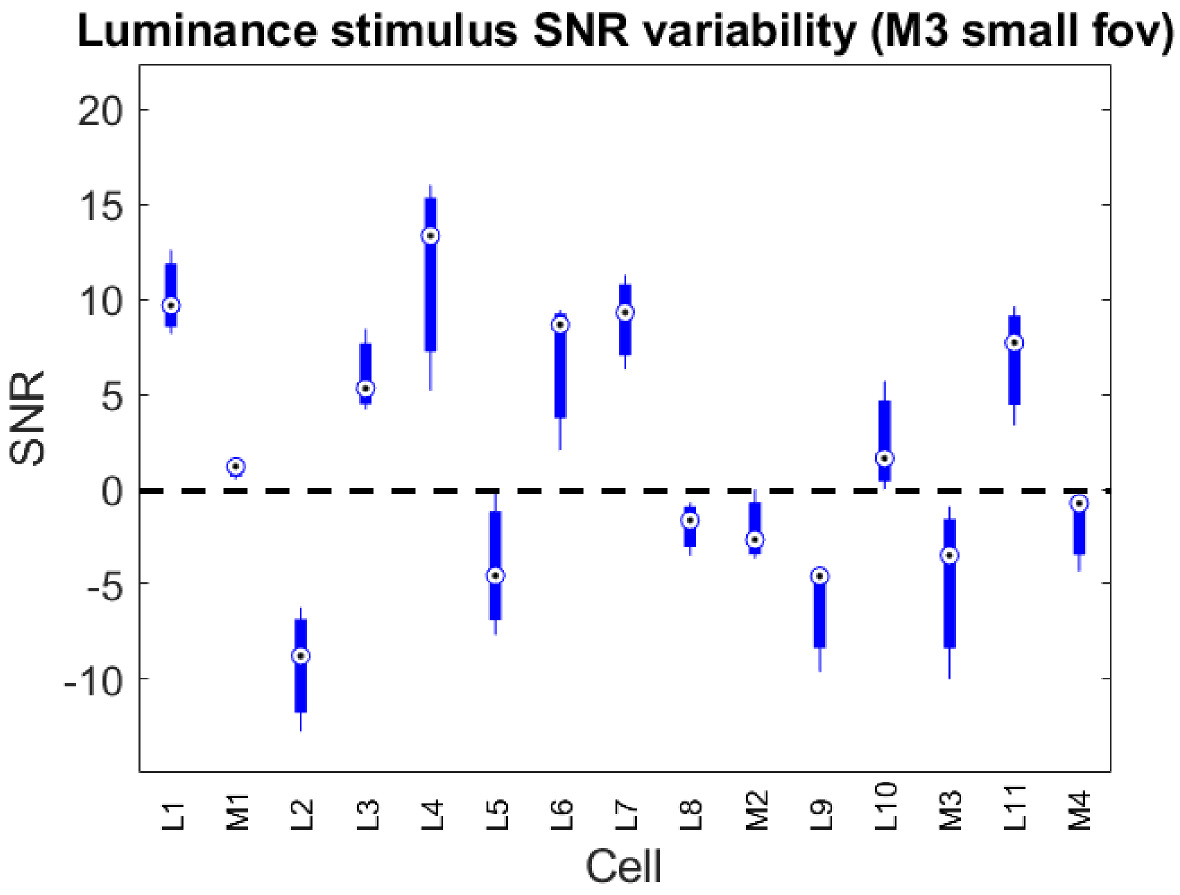
Luminance stimulus SNR variability (M3 small fov)
SNR
20
15
10
5
0
-5
-10
L1
M1
L2
L3
L4
L5
L6
L7
L8
M2
L9
L10
M3
L11
M4
Cell

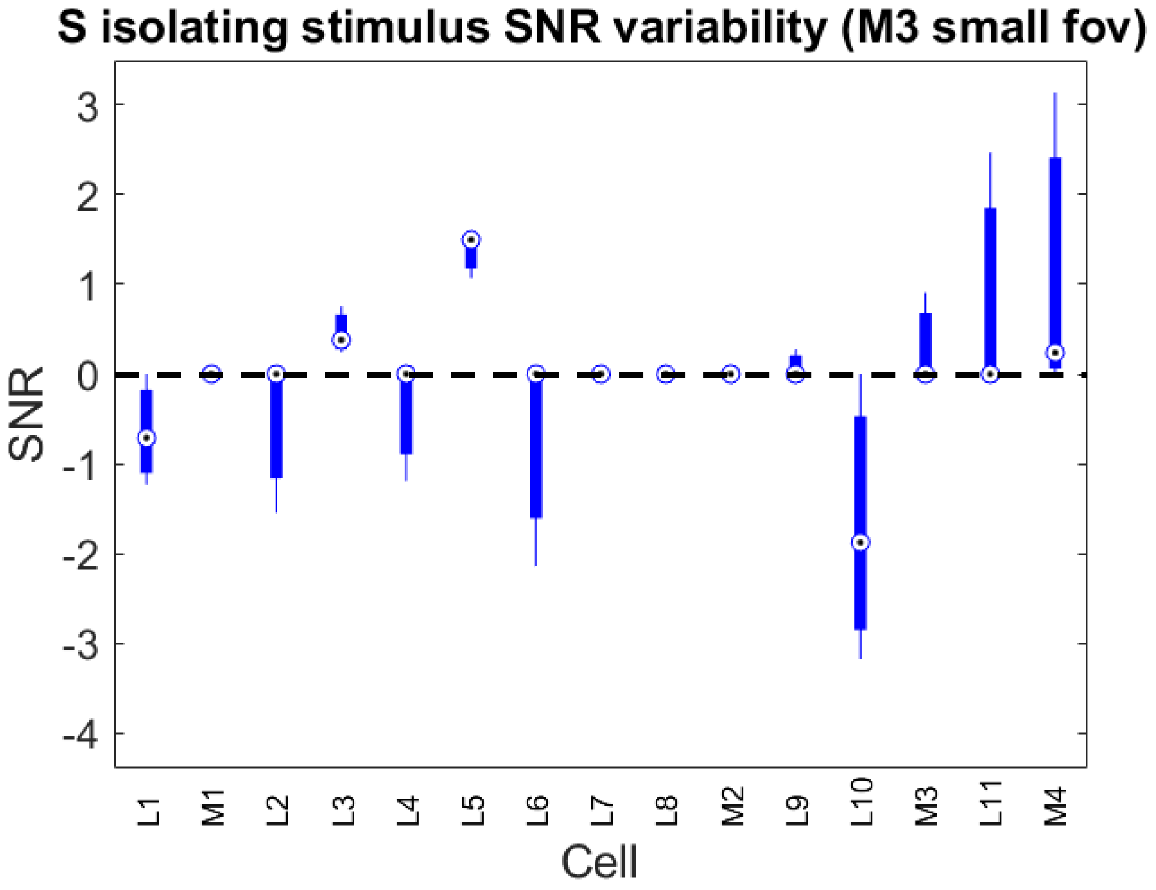
S isolating stimulus SNR variability (M3 small fov)
SNR
3
2
1
0
-1
-2
-3
-4
L1
M1
L2
L3
L4
L5
L6
L7
L8
M2
L9
L10
M3
L11
M4
Cell

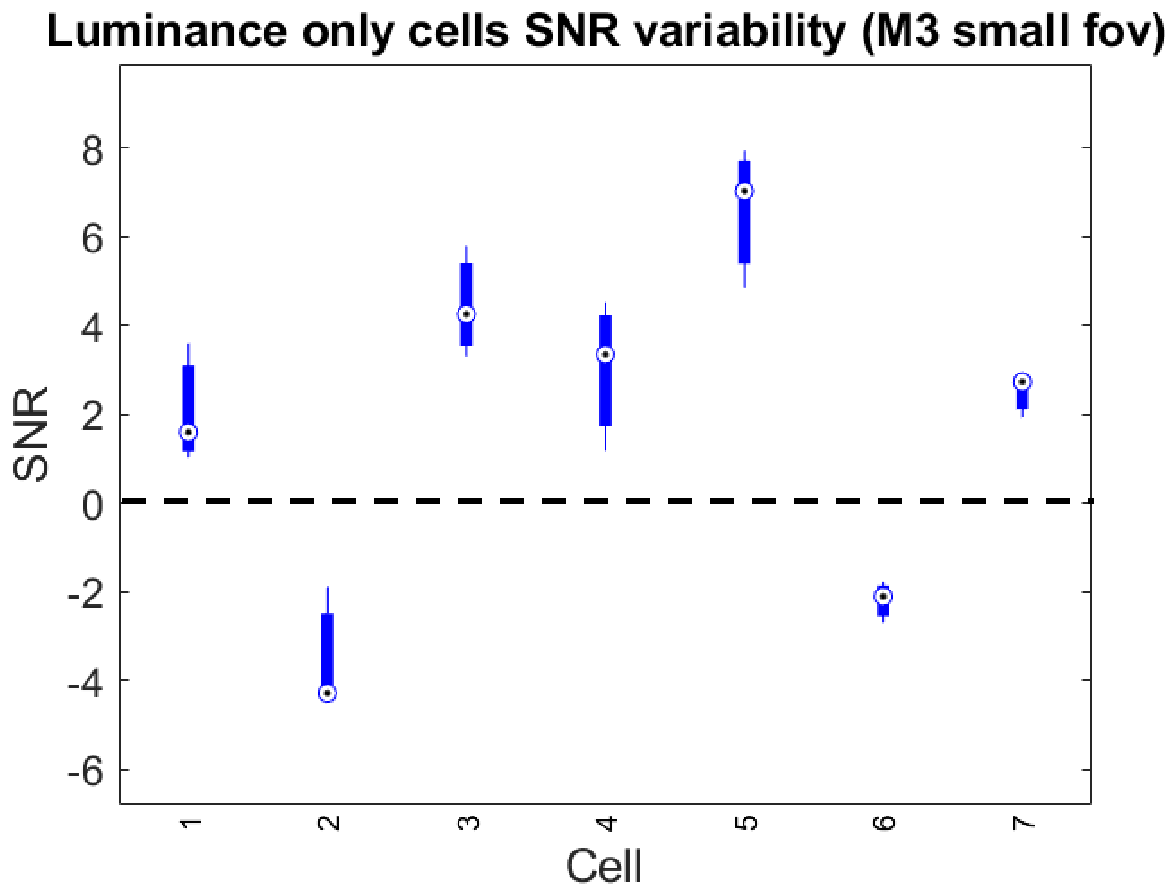

Luminance only cells SNR variability (M3 small fov)
SNR
Cell
8
6
4
2
0
-2
-4
-6
1
2
3
4
5
6
7


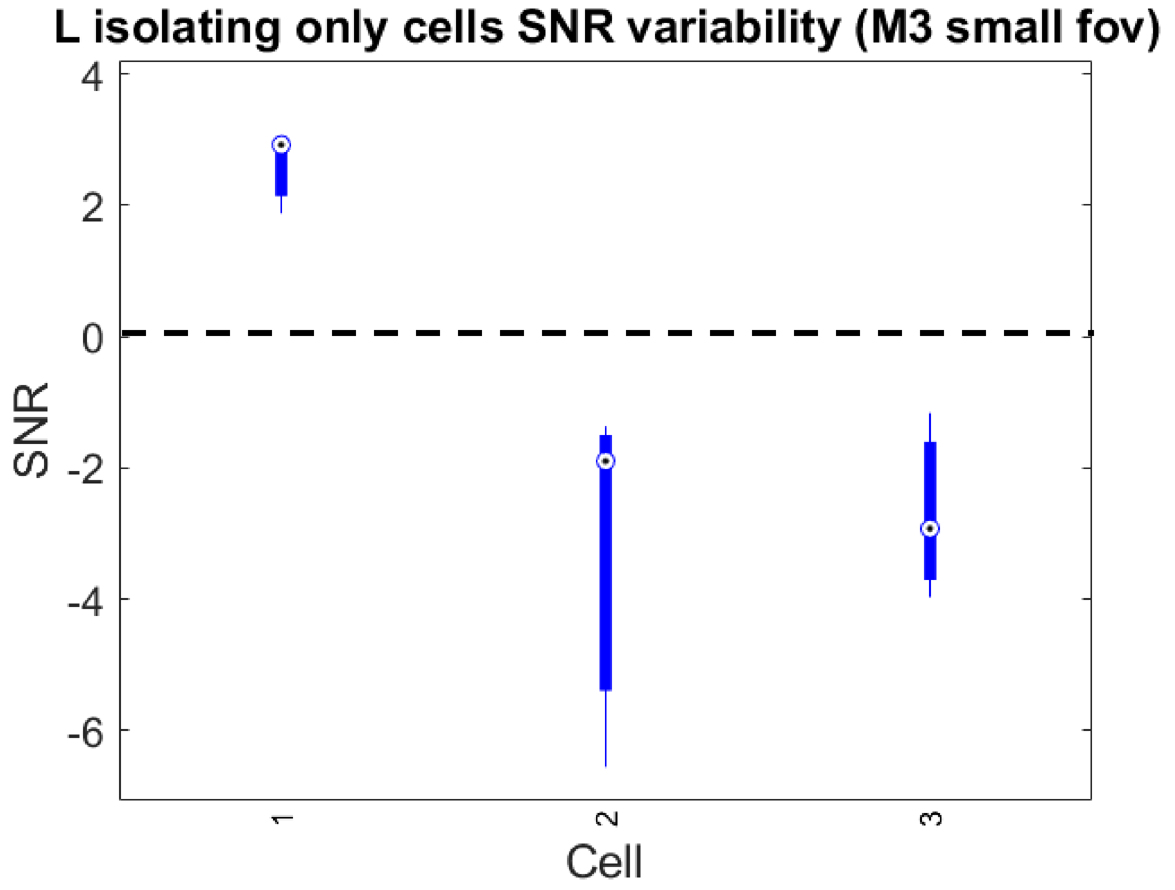

L isolating only cells SNR variability (M3 small fov)
SNR
Cell
4
2
0
-2
-4
-6
1
2
3

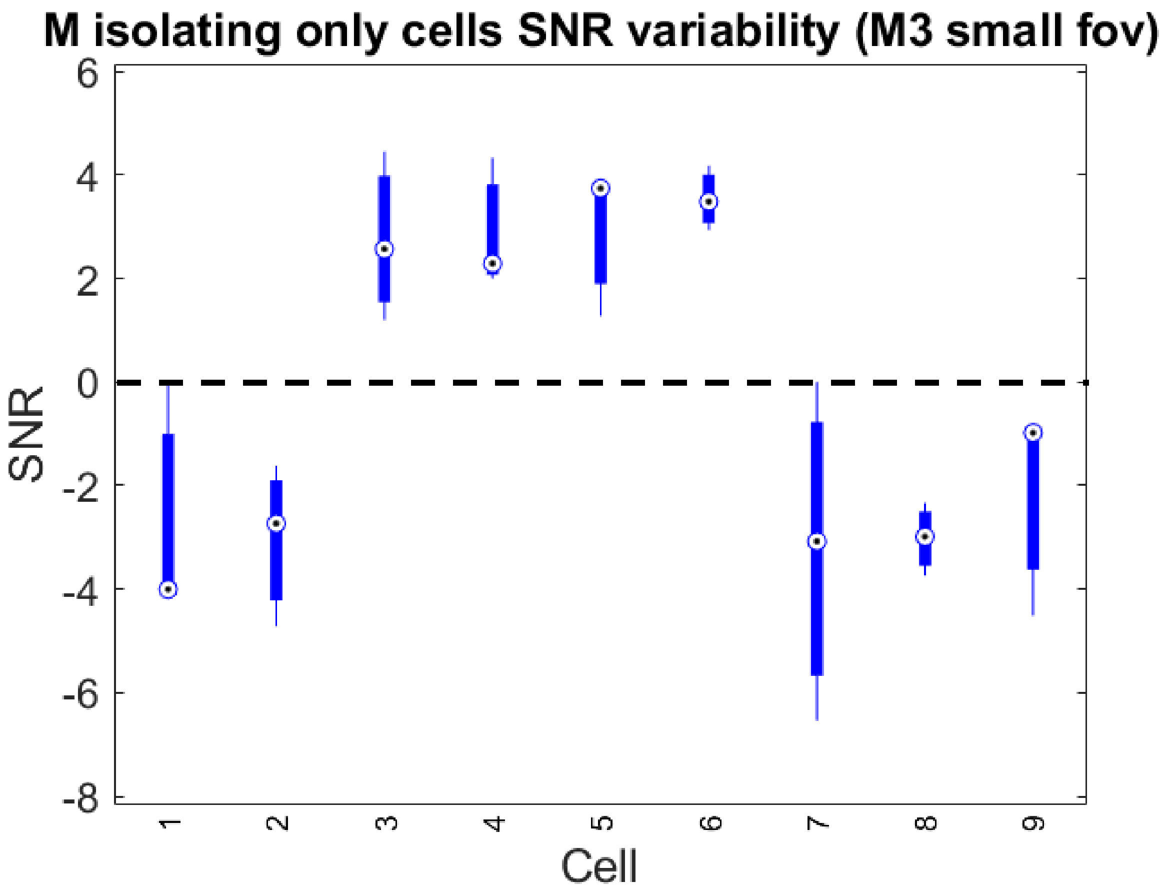

M isolating only cells SNR variability (M3 small fov)
SNR
Cell
6
4
2
0
-2
-4
-6
-8
1
2
3
4
5
6
7
8
9

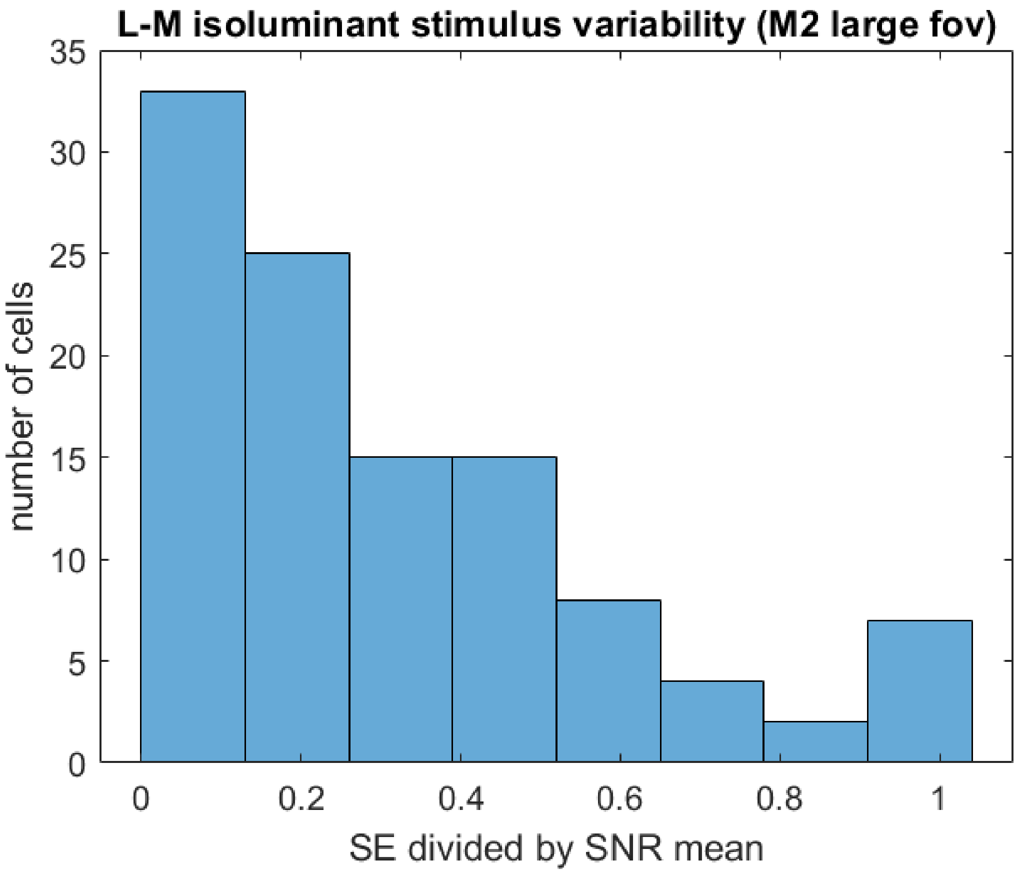

L-M isoluminant stimulus variability (M2 large fov)
35
30
25
20
15
10
5
0
number of cells
0
0.2
0.4
0.6
0.8
1
SE divided by SNR mean


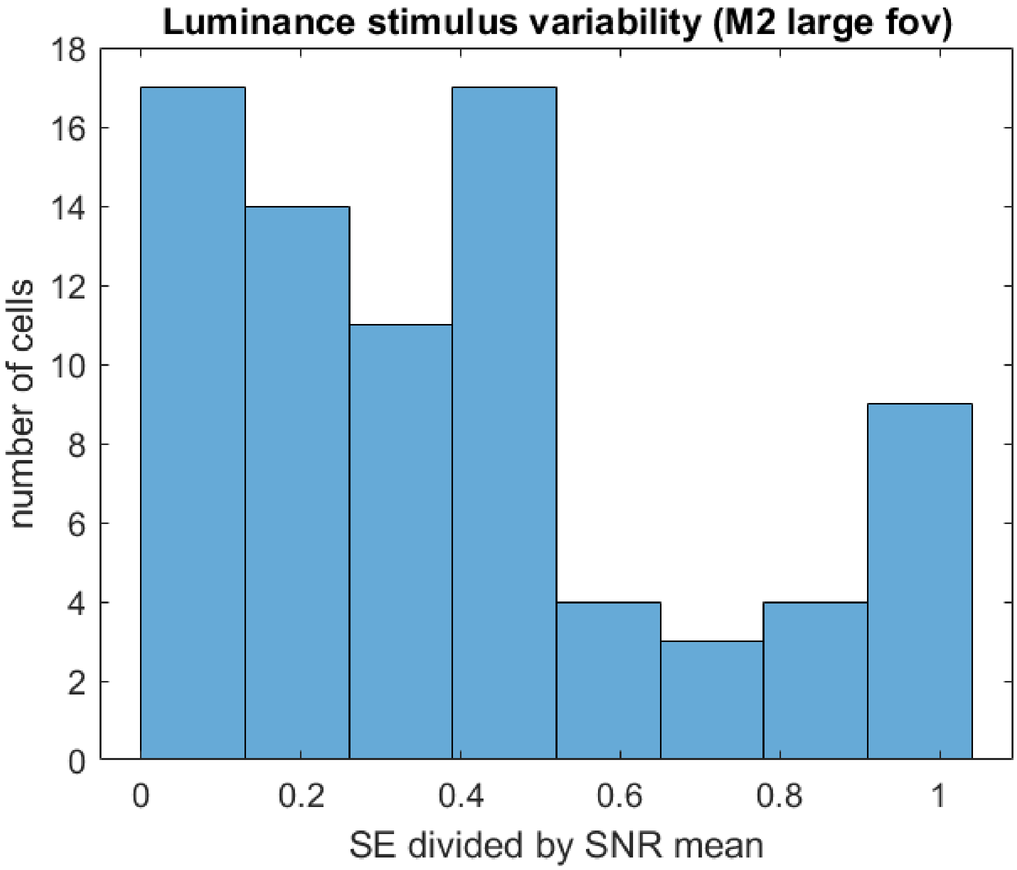

Luminance stimulus variability (M2 large fov)
18
16
14
12
10
8
6
4
2
0
number of cells
0
0.2
0.4
0.6
0.8
1
SE divided by SNR mean

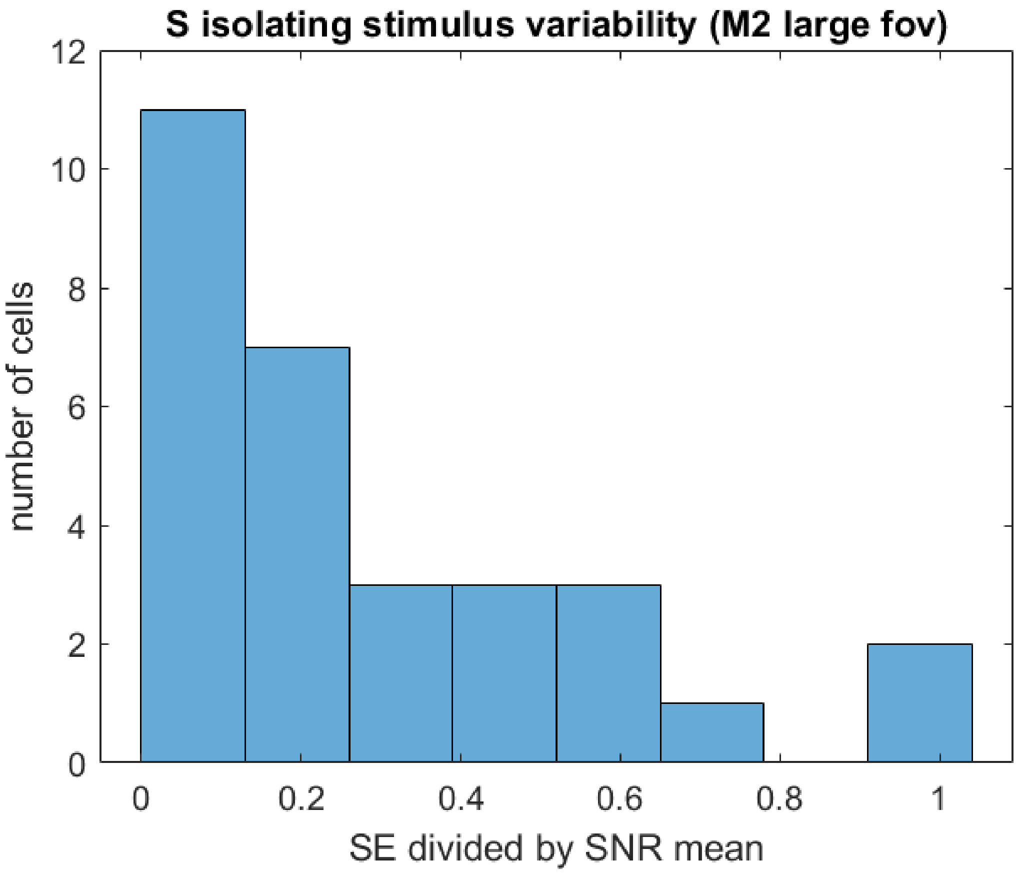
S isolating stimulus variability (M2 large fov)
number of cells
12
10
8
6
4
2
0
0
0.2
0.4
0.6
0.8
1
SE divided by SNR mean

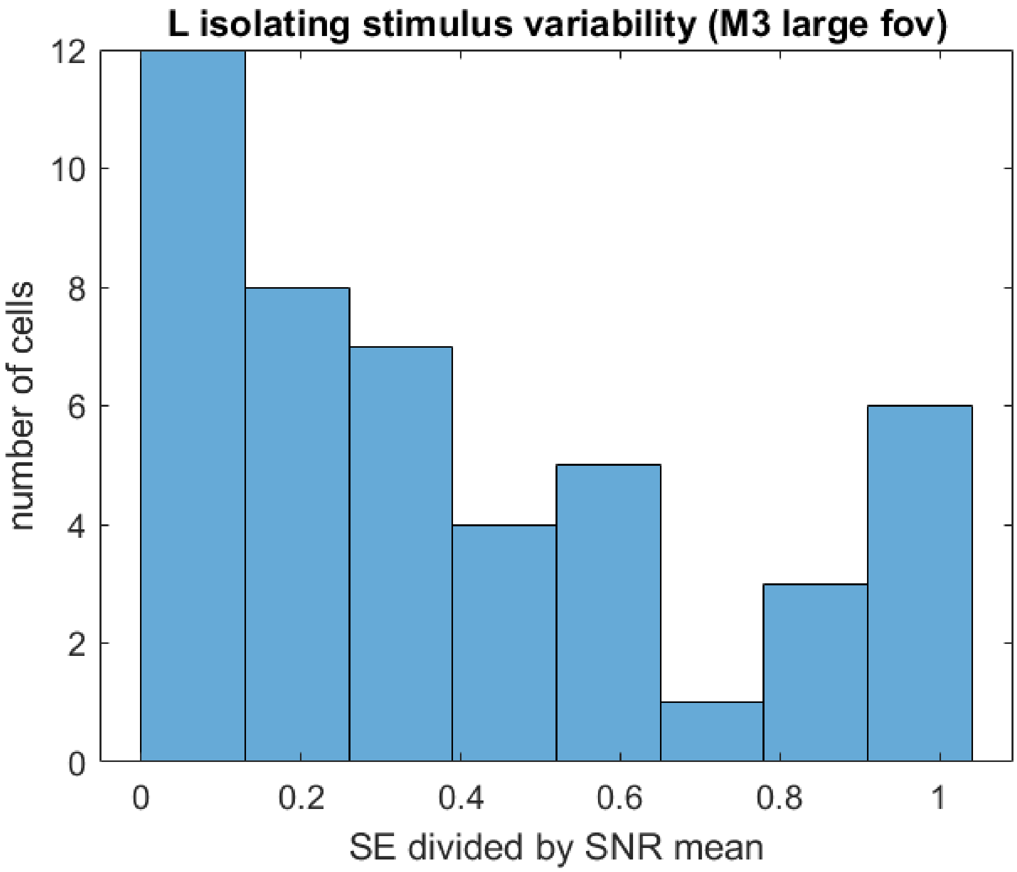
L isolating stimulus variability (M3 large fov)
number of cells
12
10
8
6
4
2
0
0
0.2
0.4
0.6
0.8
1
SE divided by SNR mean

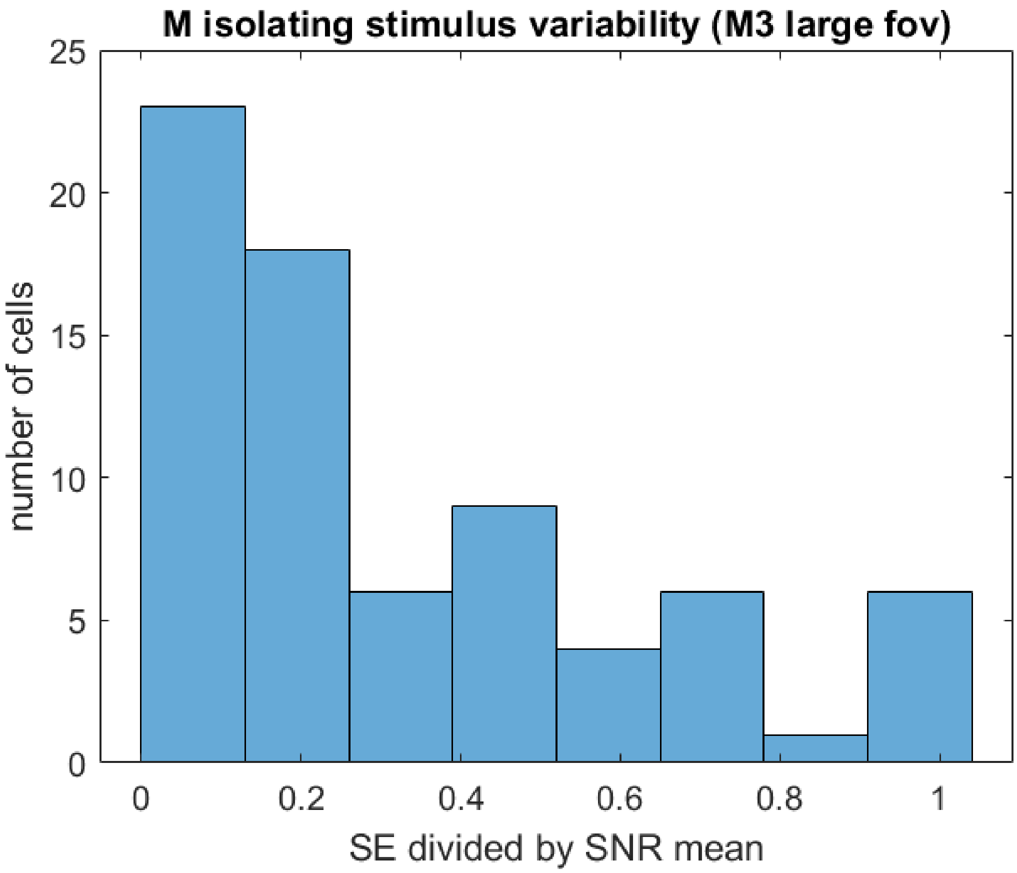
M isolating stimulus variability (M3 large fov)
number of cells
SE divided by SNR mean
0
5
10
15
20
25
0
0.2
0.4
0.6
0.8
1

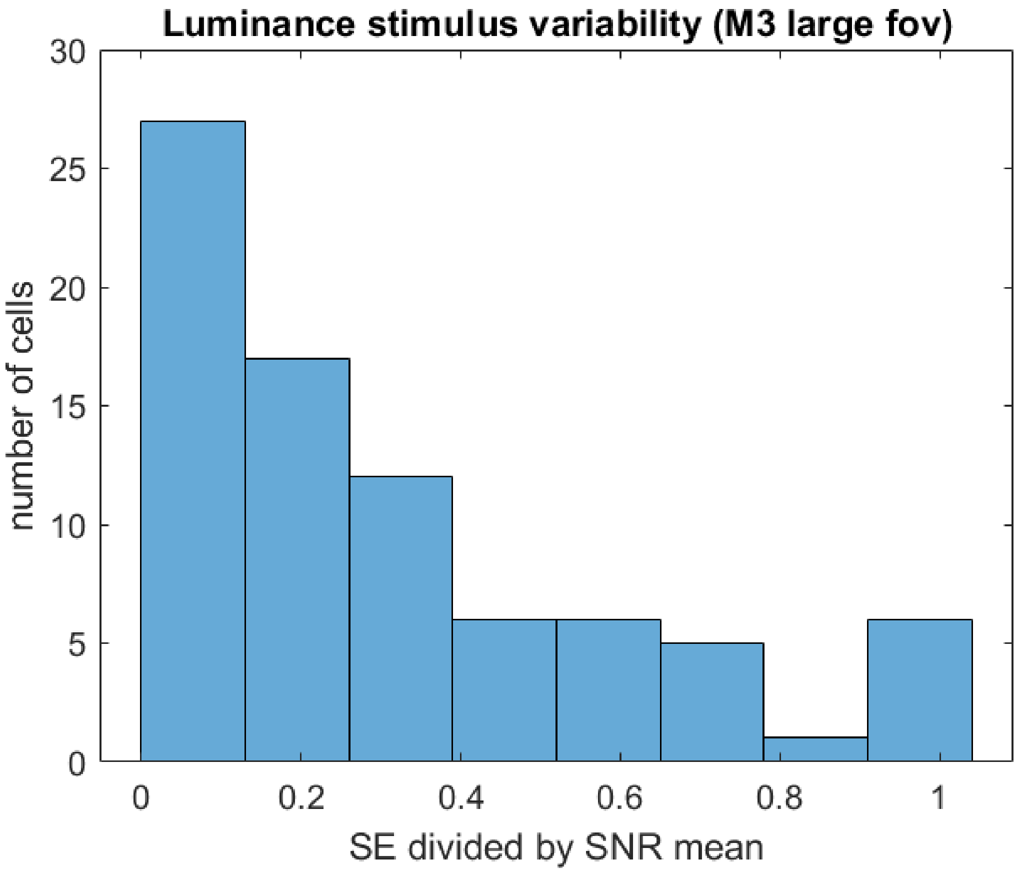
Luminance stimulus variability (M3 large fov)
number of cells
SE divided by SNR mean
0
5
10
15
20
25
30
0
0.2
0.4
0.6
0.8
1

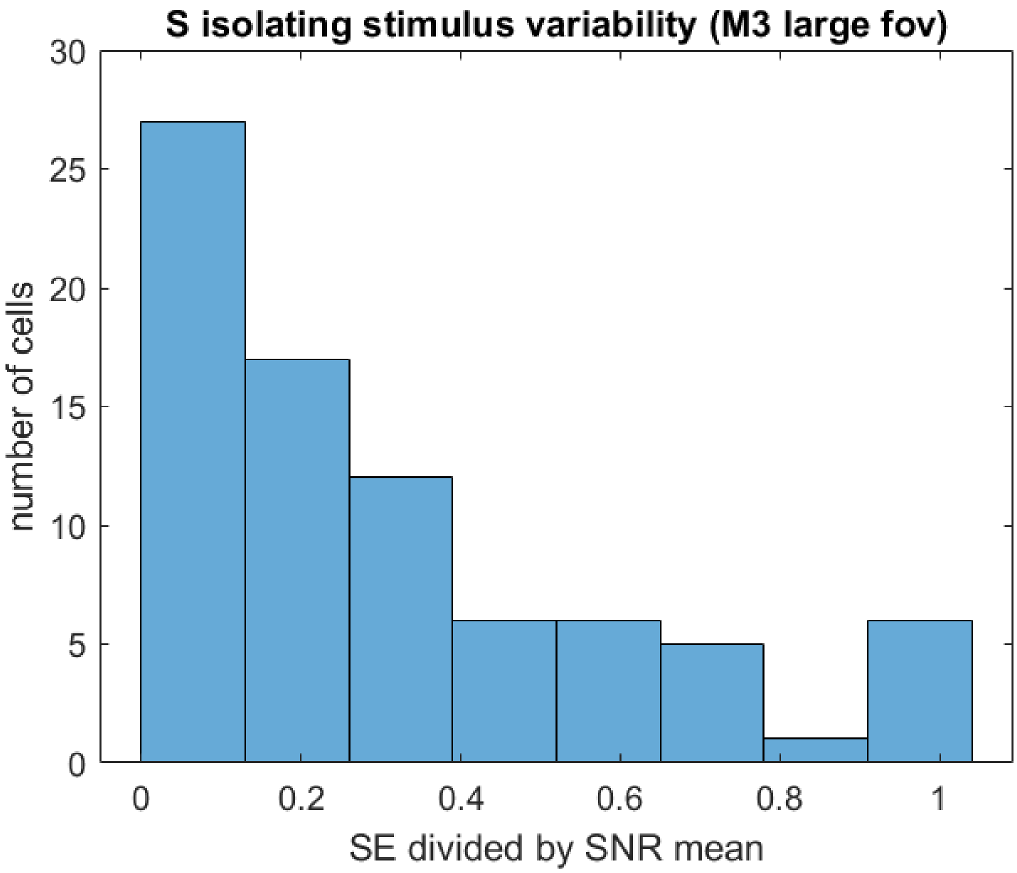


**S3 Fig. Variability of responses to chromatic and achromatic flicker.** Plots for the small FOV from M3 show the response variability to the L isolating, M isolating, S isolating, and Luminance stimuli. Note that these are boxplots showing the median, 25-75% interquartile range, and lowest and highest values, but there were only three measurements. Thus, the median value and the two ends of the boxplots represent the three measured values for each cell. For the large FOV from M3 and M2, there are many more cells, so the standard error divided by the SNR mean is plotted as a histogram for all cells that were responsive to each individual stimulus. No standard error can be plotted for M1, as there was only one experiment in that animal.
