## Supplement 4 for "*In vivo* chromatic and spatial tuning of foveolar retinal ganglion cells in *Macaca fascicularis*"

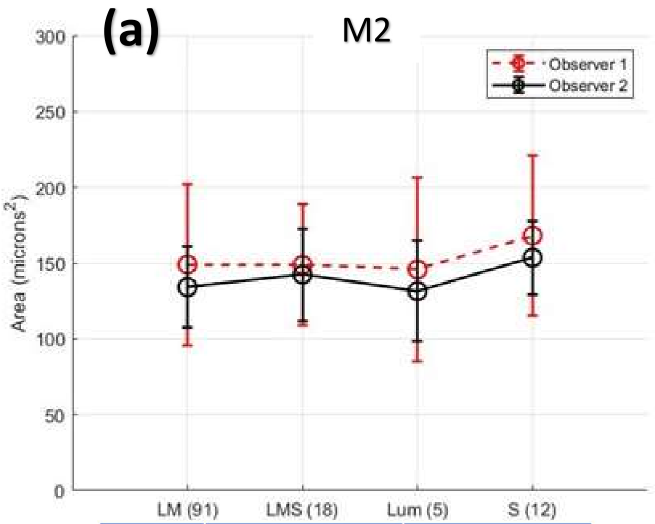


| | Mean | SD |
|---|---|---|
| Observer 1 | | |
| L-M | 149.0679 | 53.3055 |
| L-M/S | 148.8017 | 39.9526 |
| LUM | 145.8026 | 60.7404 |
| S | 168.2190 | 52.6120 |
| Observer 2 | | |
| L-M | 134.1611 | 26.9083 |
| L-M/S | 142.3934 | 30.3989 |
| LUM | 131.4992 | 33.2721 |
| S | 153.6080 | 24.1251 |

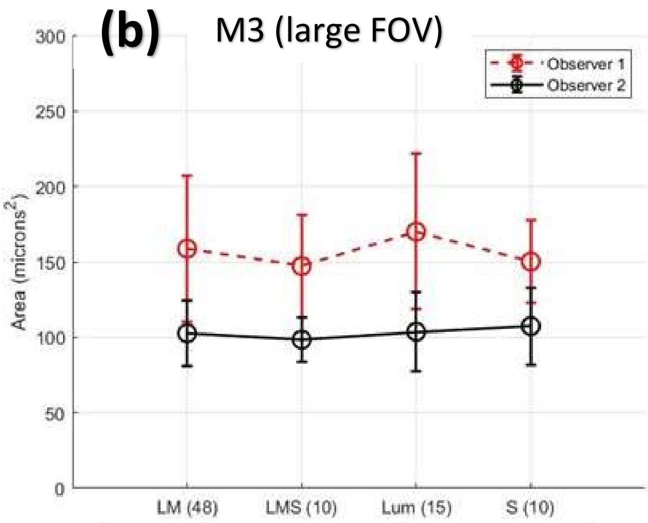


| | Mean | SD |
|---|---|---|
| Observer 1 | | |
| L-M | 158.8273 | 48.3035 |
| L-M/S | 147.2437 | 33.9356 |
| LUM | 170.0995 | 51.4246 |
| S | 150.3205 | 27.0788 |
| Observer 2 | | |
| L-M | 102.7410 | 21.9930 |
| L-M/S | 98.4555 | 14.8620 |
| LUM | 103.7299 | 26.2566 |
| S | 107.4660 | 25.6175 |

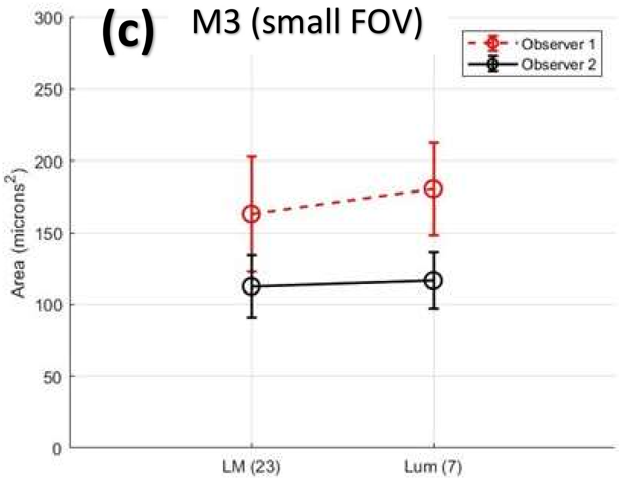


| | Mean | SD |
|---|---|---|
| Observer 1 | | |
| L-M | 162.9411 | 39.6726 |
| LUM | 180.5915 | 32.4137 |
| Observer 2 | | |
| L-M | 112.5965 | 21.5090 |
| LUM | 116.6258 | 19.8030 |

**(d)**

| M2 Large FOV – Observer 1 | | | | |
|---|---|---|---|---|
| | L-M | L-M/S | Lum | S |
| L-M | - | 0.7194 | 0.7105 | 0.1442 |
| L-M/S | 0.7194 | - | 0.7093 | 0.2897 |
| Lum | 0.7105 | 0.7093 | - | 0.4421 |
| S | 0.1442 | 0.2897 | 0.4421 | - |

| M2 Large FOV – Observer 2 | | | | |
|---|---|---|---|---|
| | L-M | L-M/S | Lum | S |
| L-M | - | 0.2572 | 0.6723 | 0.0118 |
| L-M/S | 0.2572 | - | 0.4103 | 0.2777 |
| Lum | 0.6723 | 0.4103 | - | 0.1354 |
| S | 0.0118 | 0.2777 | 0.1354 | - |

| M3 Large FOV – Observer 1 | | | | |
|---|---|---|---|---|
| | LM | LMS | Lum | S |
| L-M | - | 0.6356 | 0.2167 | 1 |
| L-M/S | 0.6356 | - | 0.1827 | 0.6495 |
| Lum | 0.2167 | 0.1827 | - | 0.2218 |
| S | 1 | 0.6495 | 0.2218 | - |

| M3 Large FOV – Observer 2 | | | | |
|---|---|---|---|---|
| | L-M | L-MS | Lum | S |
| L-M | - | 0.7913 | 0.9475 | 0.5230 |
| L-M/S | 0.7913 | - | 1 | 0.4565 |
| Lum | 0.9475 | 1 | - | 0.6121 |
| S | 0.5230 | 0.4565 | 0.6121 | - |

| M3 Small FOV – Observer 1 | |
|---|---|
| | L-M |
| Lum | 0.2915 |

| M3 Small FOV – Observer 2 | |
|---|---|
| | L-M |
| Lum | 0.6404 |

**S4 Fig. Soma sizes of RGCs for animals M2 and M3 across chromatic functional groups.** Observer 1 used the open source software GIMP to segment RGCs in fluorescence images from M2 (large FOV), M3 (large FOV) and M3 (small FOV). The ellipse tool was used to segment the rough boundary of individual RGCs. Observer 2 used the open source software ImageJ to segment the same RGCs in fluorescence images from M2 and M3 using a hand tracing tool to trace the observable edges of each cell's fluorescence. Under both methods, the area for each cell was computed in terms of pixels$^2$ and then converted to μm$^2$ using the following formula: $\frac{\mu m}{pixel} = 291.2\ \frac{\mu m}{deg} * \frac{axial\ length}{24.2\ mm} * \frac{FOV\ width}{496\ pixels}$, where 291.2 μm/deg is the human model eye visual angle to retinal extent conversion, 24.2 mm is the human model eye axial length, 496 pixels is the width of the imaging PMT used in the AOSLO system, axial length is the animal's axial length in mm, and FOV width is the width in degrees of the FOV used. The axial length of animal M2 is 17.2 mm, and the FOV width used was 3.64 deg. The axial length of animal M3 is 16.56 mm, and the FOV widths used were 3.69 deg (large FOV) and 2.54 deg (small FOV). Cells were compared across functional groups identified as L-M/M-L chromatic opponent (L-M), S only responding (S), Luminance only or achromatic (LUM), and mixed L-M/S responses (L-M/S). In **(a)**, the soma areas in μm$^2$ for the four functional groups in M2 are listed as the mean and standard deviation of each group across both observers. In **(b)**, the same comparisons are made for the four functional groups in M3 at the large FOV. In **(c)**, comparisons for the two functional groups found in M3 at the inner edge of the foveal slope (small FOV) are made across both observers. In, **(d)**, summary tables show the p-scores from a Mann-Whitney U test (MATLAB function ranksum(x,y)) comparing the distributions of soma areas across functional groups as measured by both observers separately. All p-values were greater than 0.1, except for Observer 2's comparison of the L-M group to the S only group in M2 which had p = 0.0118. Based on these results, we cannot reject the hypothesis that the distributions of soma sizes for these functional groups are roughly the same across the two animals measured at the range of eccentricities at which we imaged cell somas.
