## Supplement 5 for "*In vivo* chromatic and spatial tuning of foveolar retinal ganglion cells in *Macaca fascicularis*"

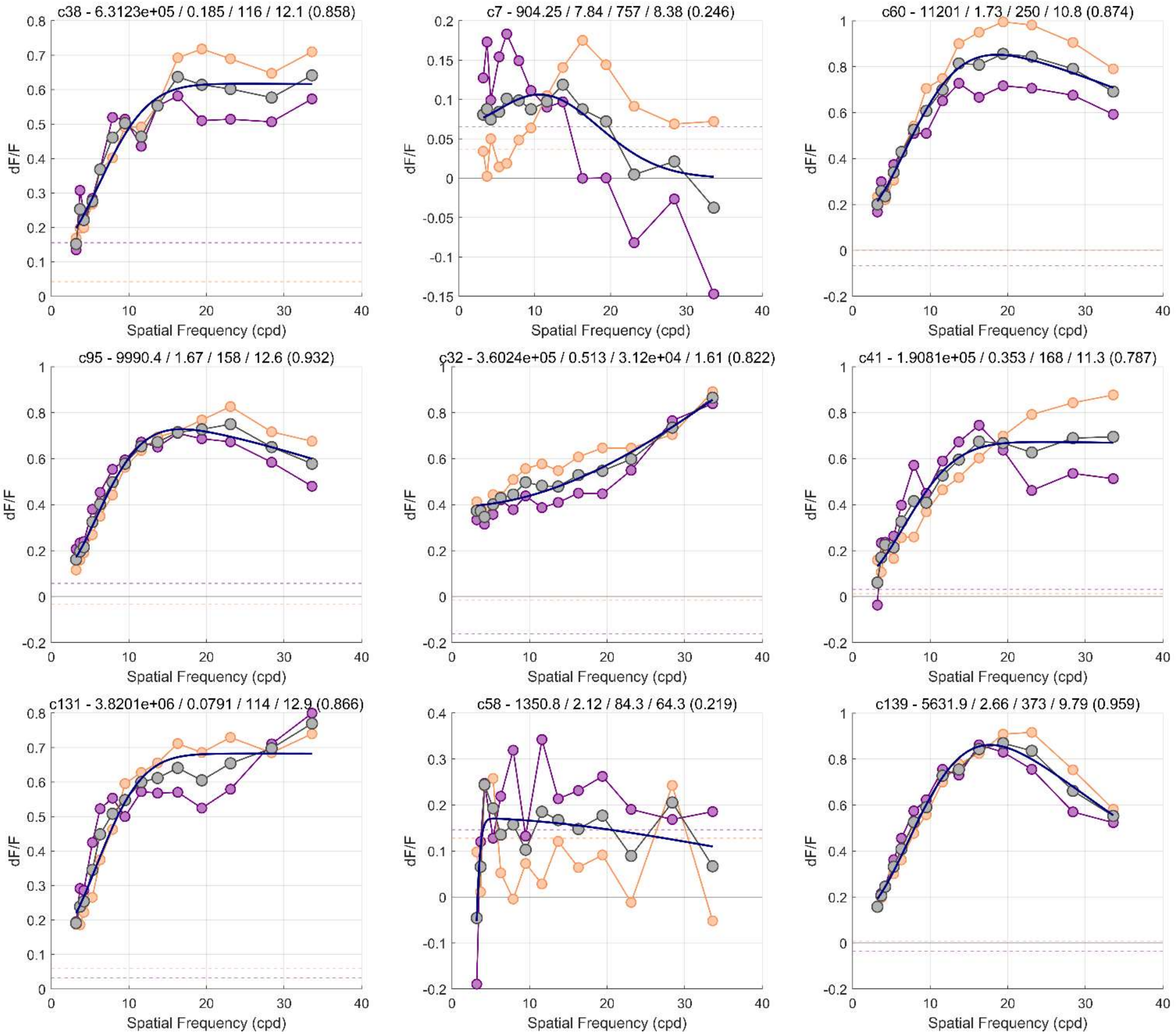


**S5 Fig. Spatial response of RGCs from M3 in the large FOV condition.** Some example responses of L-M/M-L chromatic opponent RGCs from M3 at the large FOV, where each plot is the response of a different cell to drifting gratings (6 Hz) of spatial frequencies from 2 – 34 c/deg. These cells were chosen to showcase different cell responses and are not necessarily special or representative of the population. Purple and Orange curves show two separate experiments that were averaged together (Gray curve). The Blue curves are simple difference of Gaussians fits (not the full ISETBio modeling, for reasons explained below) to each average. Each plot title includes the cell's unique label "cN", and five parameters—the center strength $K_c$, the center radius $r_c$, the surround strength $K_s$, the surround radius $r_s$, and a goodness of fit value for the difference of Gaussians. As can be easily seen, many of the cells do not exhibit high spatial frequency falloff at the range of spatial frequencies measured, and so the difference of Gaussians either fits poorly or produces fit parameters that are highly irregular or suspect such as center sizes much smaller than the size of a single cone. For this reason, these data were not incorporated into the modelling in the main body, as higher spatial frequency responses were needed to properly fit many of these cell responses. There were 48 such L-M/M-L chromatic opponent cells in the large FOV in M3 out of 145 measured cells. In M2, there was only one experiment measuring response to spatial frequency and it suffered from the same lack of high spatial frequency measurements as the data shown here. Full data from both animals can be shared upon request.
