## Supplement 6 for "*In vivo* chromatic and spatial tuning of foveolar retinal ganglion cells in *Macaca fascicularis*"

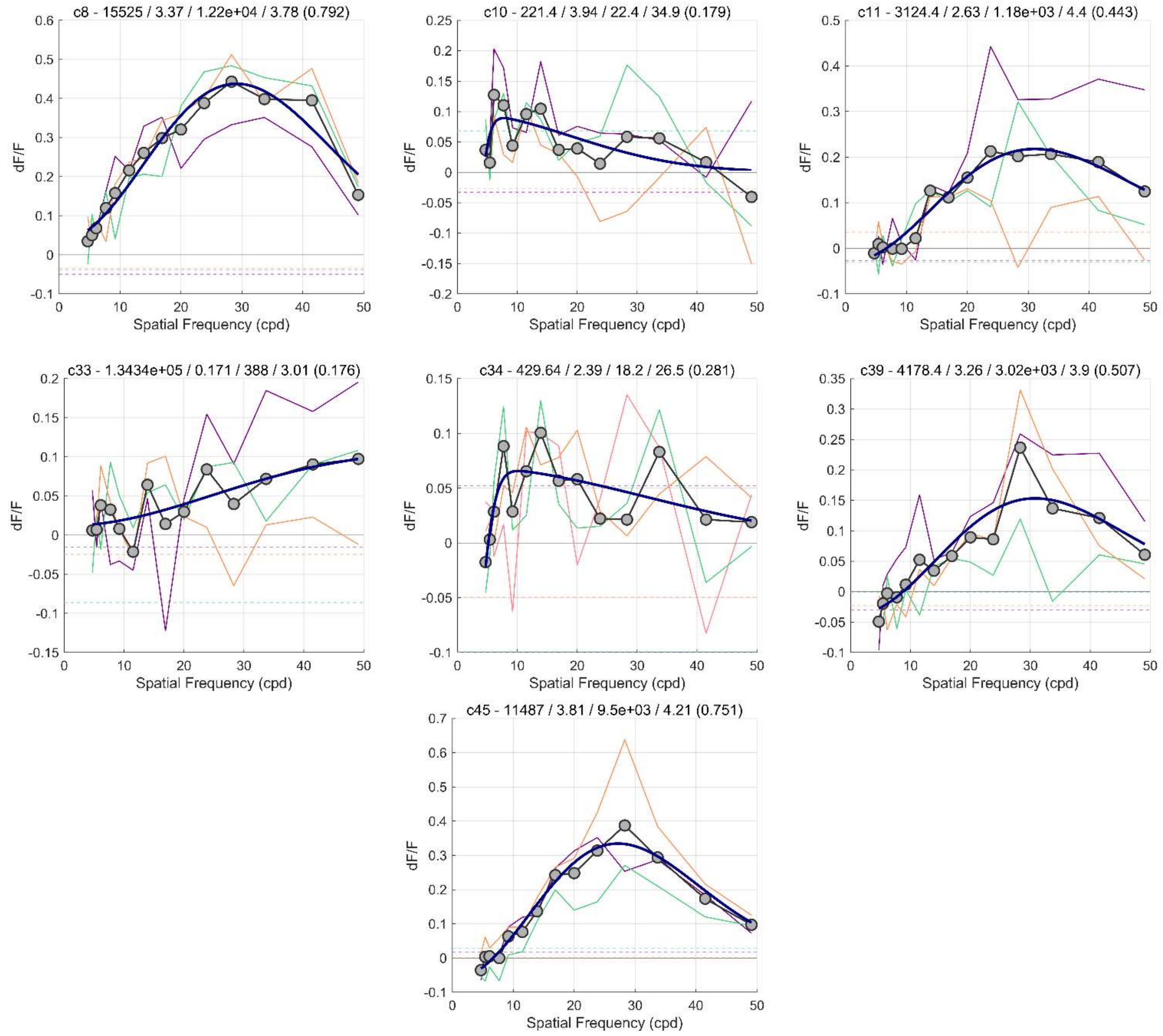


**S6 Fig. Spatial response of achromatic RGCs from M3 at the center of the foveola.** Each graph represents a single cell identified as responding only to achromatic stimuli from the center of the foveola of M3. The measured responses were to drifting gratings (6 Hz) of spatial frequencies from 4 – 49 c/deg. Purple, Orange, and Green lines represent the cell's response in three different experiments. The Gray line is the average of the three experiments for each cell, and the Blue line is a simple difference of Gaussians fit to the average (not the full ISETBio model, see S2 Fig). Each plot title includes the cell's unique label "cN", and five parameters—the center strength $K_c$, the center radius $r_c$, the surround strength $K_s$, the surround radius $r_s$, and a goodness of fit value for the difference of Gaussians. Compared to the 15 L-M and M-L cone opponent cells from this same location, these 7 achromatic cells were had noisier responses across experiments and more varied response characteristics. Though most of these cells are likely to be parasol RGCs, some could be achromatic midget RGCs or other rarer achromatic RGC types. Due to this uncertainty, these data were not included in the modelling for putative midget RGCs presented in the main manuscript. There were also achromatic cells identified in M2 and in M3 at the large FOV condition, but those data suffer from decreased resolution and a decreased range of spatial frequencies presented to the cells. Full data from all animals can be shared upon request.
