## Supplementary figures and images for "*In vivo* chromatic and spatial tuning of foveolar retinal ganglion cells in *Macaca fascicularis*"

### Supplement 7

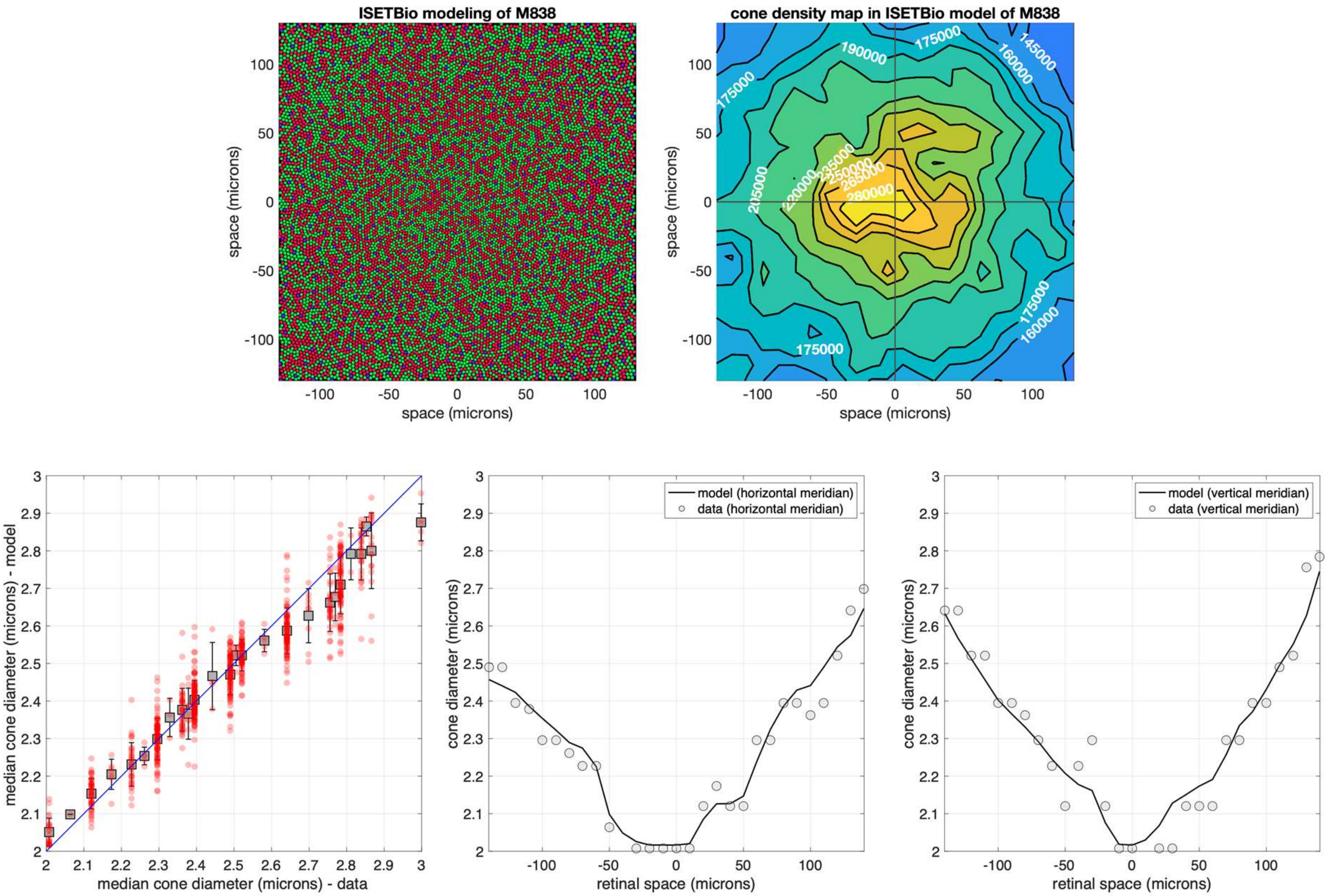

### Supplement 8

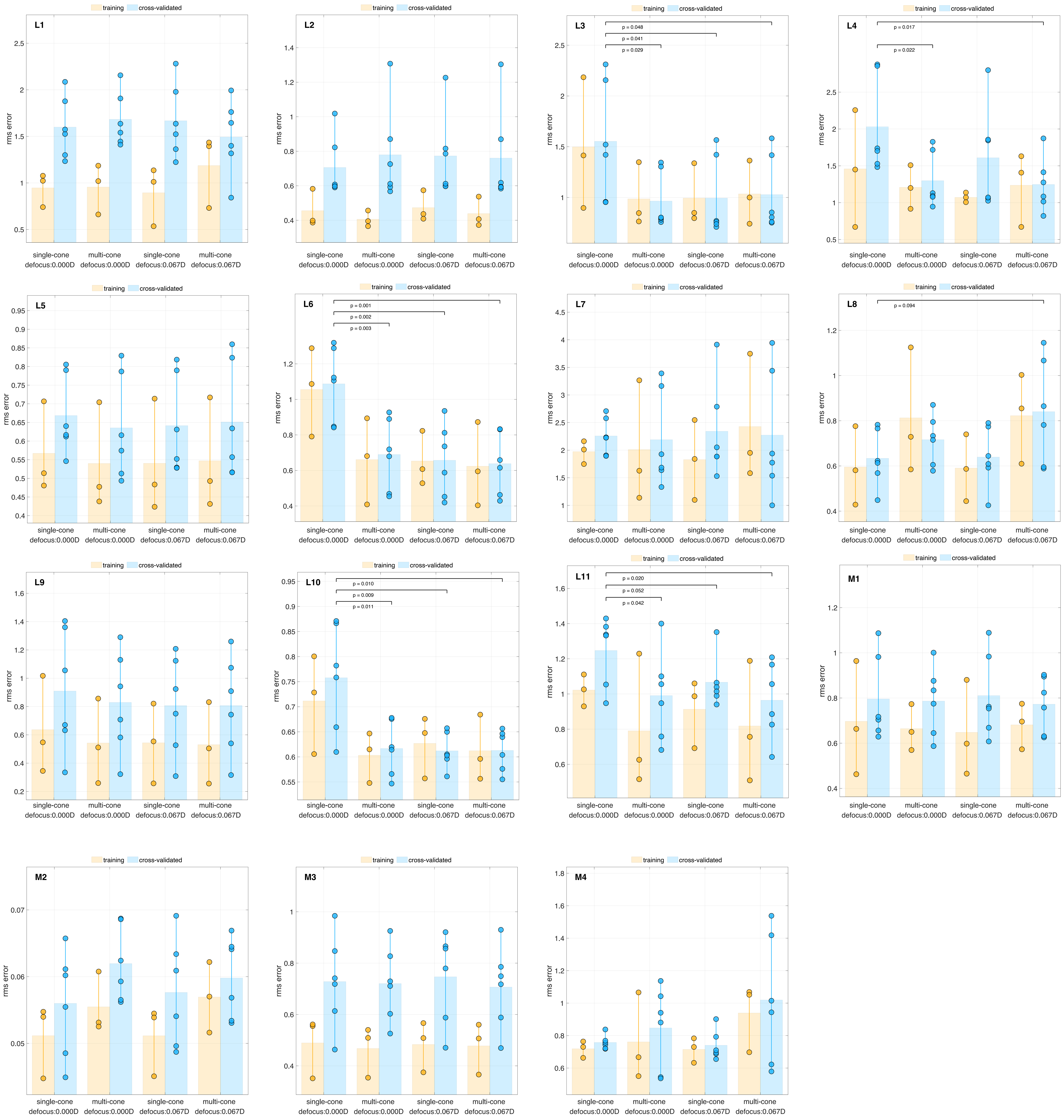

### Supplement 9

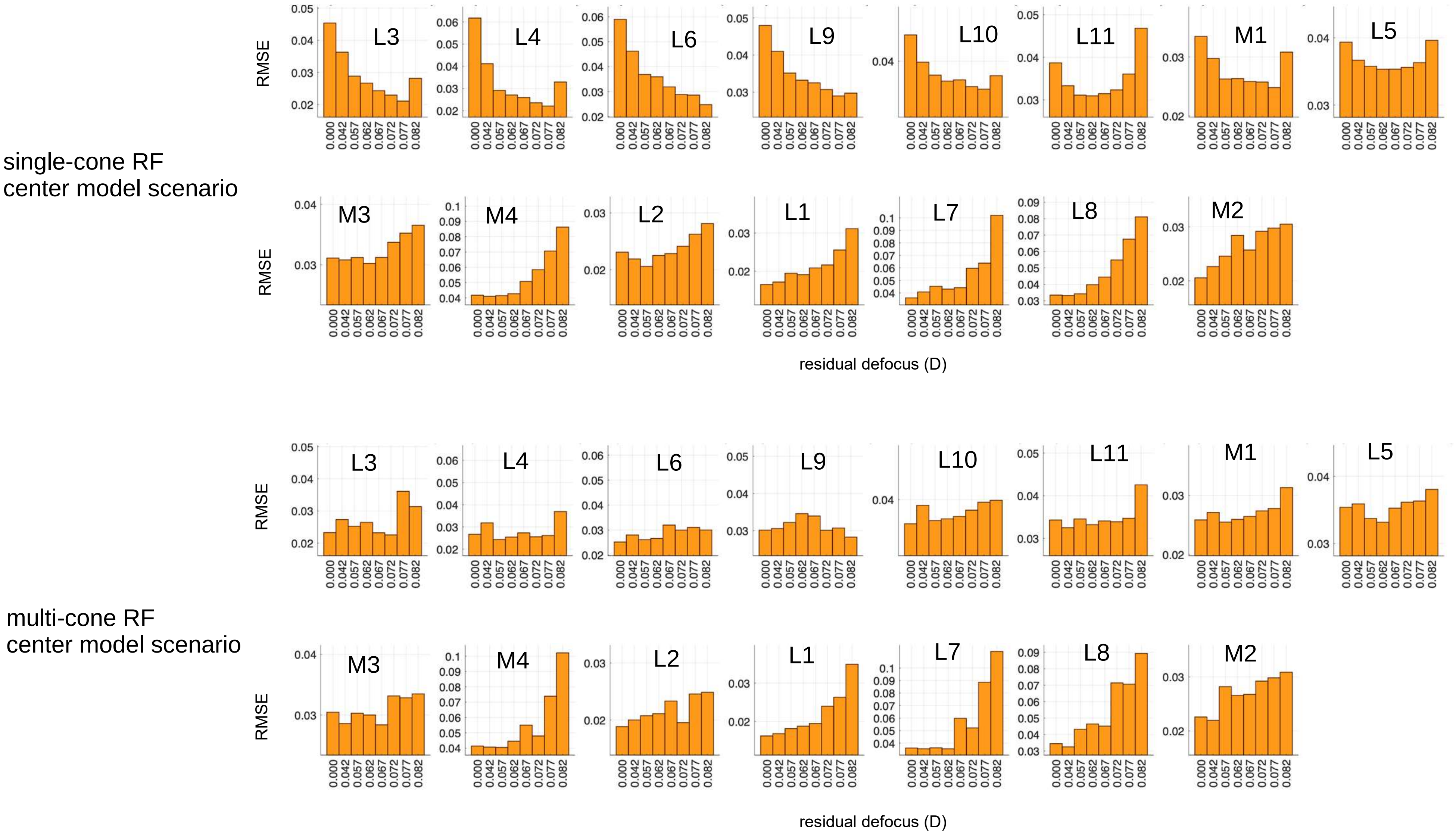
