## Supplement 10 for "*In vivo* chromatic and spatial tuning of foveolar retinal ganglion cells in *Macaca fascicularis*"

### single-cone center, 0.00D residual defocus

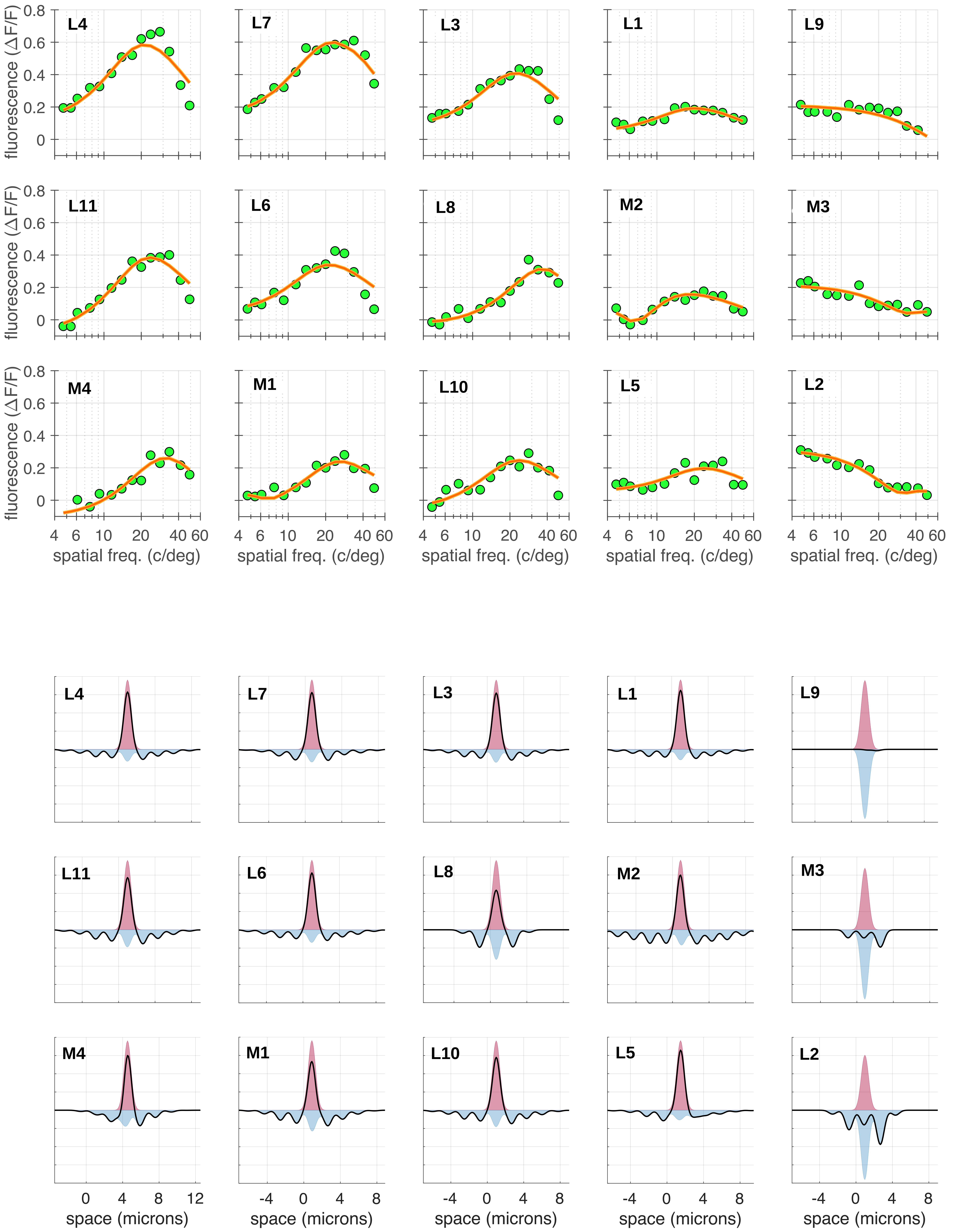

multi-cone center, 0.00D residual defocus

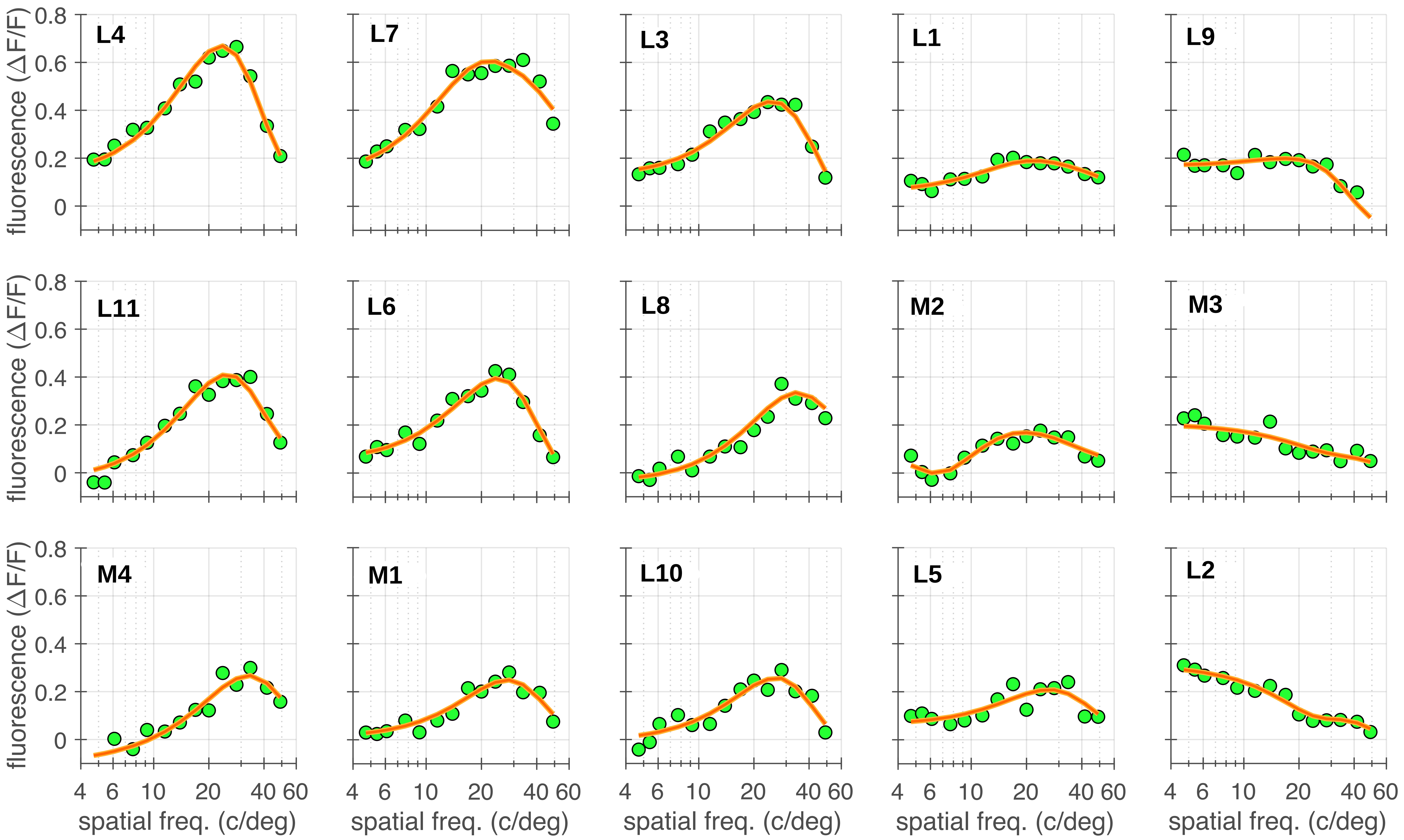


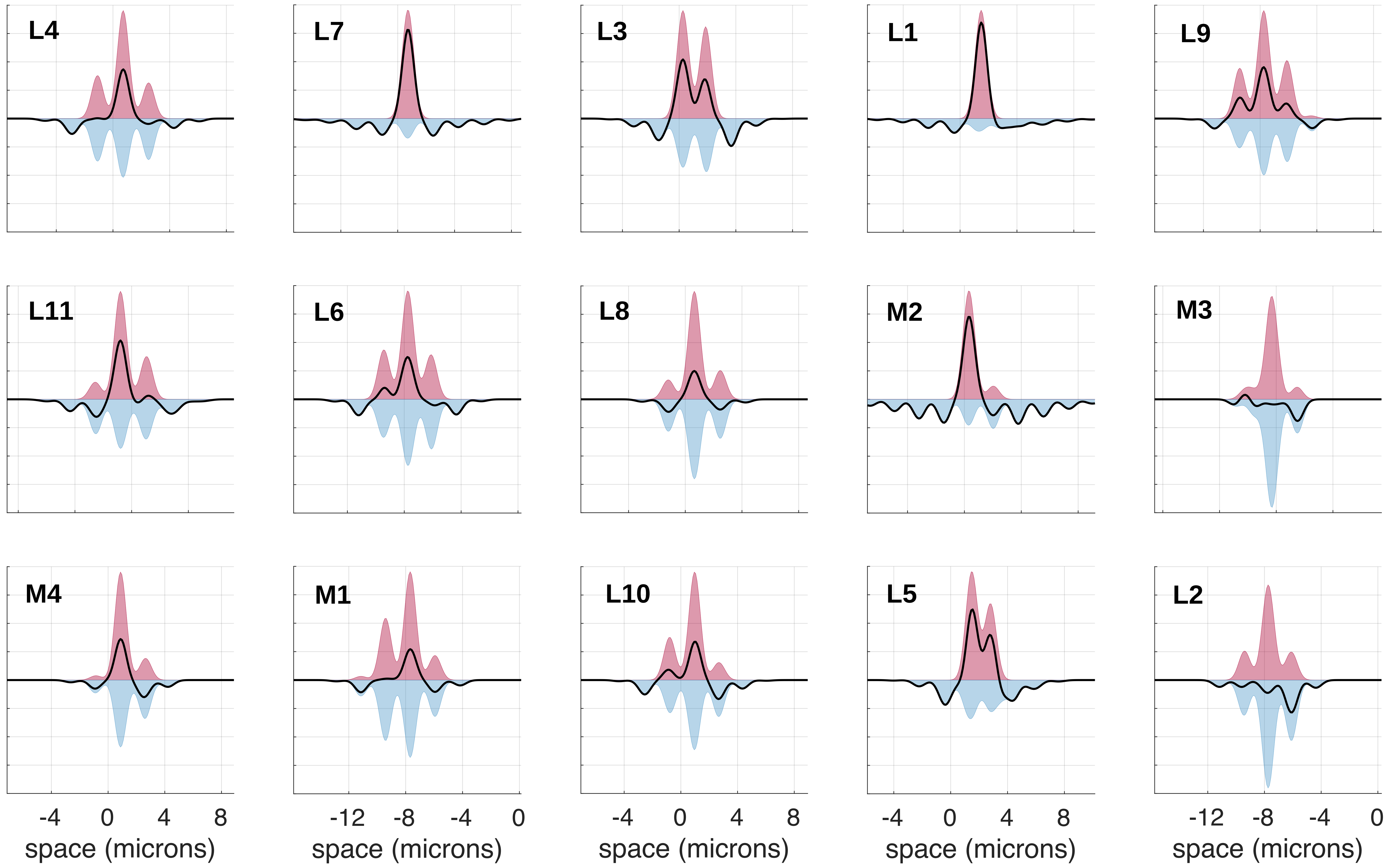

### single-cone center, 0.067D residual defocus

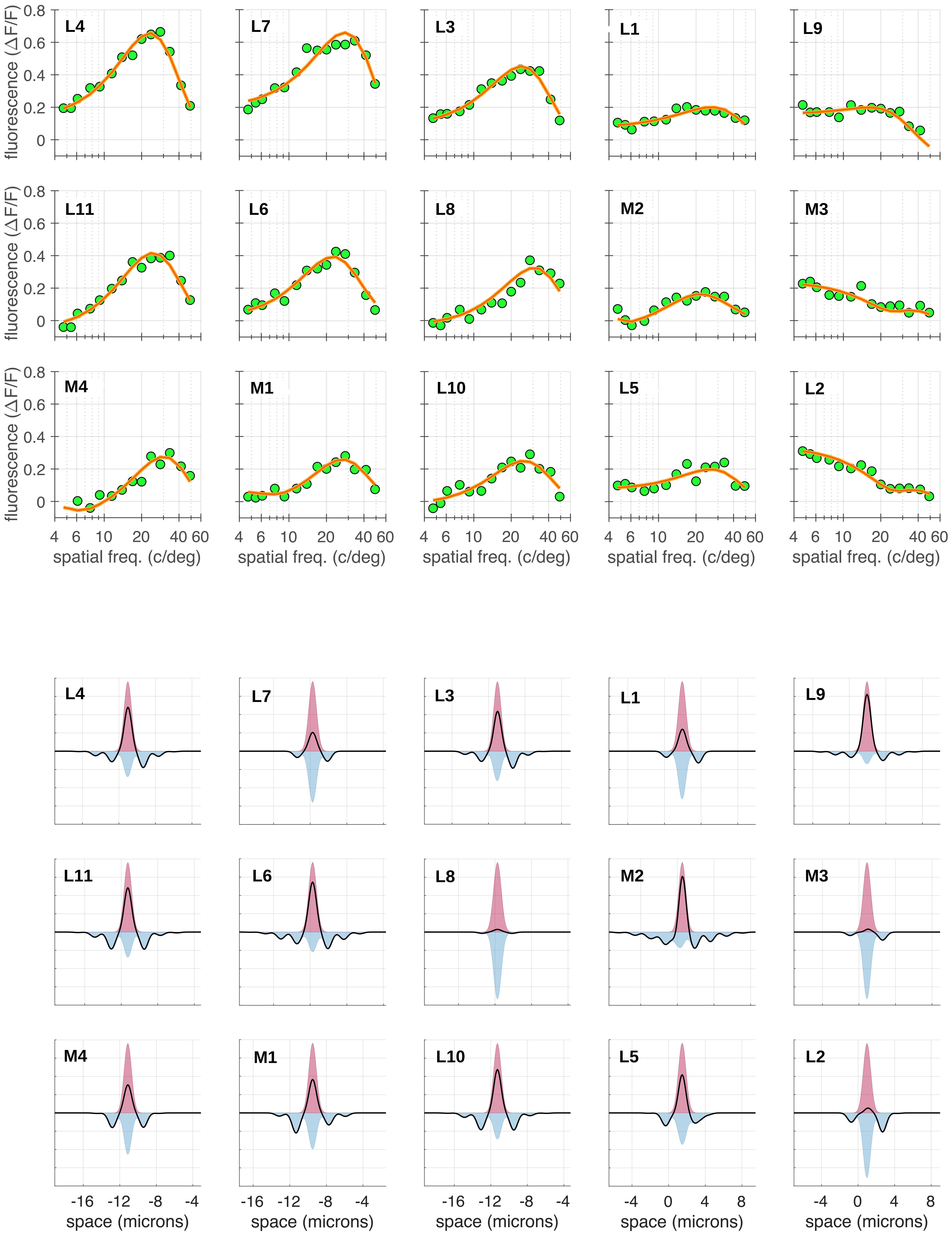

### multi-cone center, 0.067D residual defocus

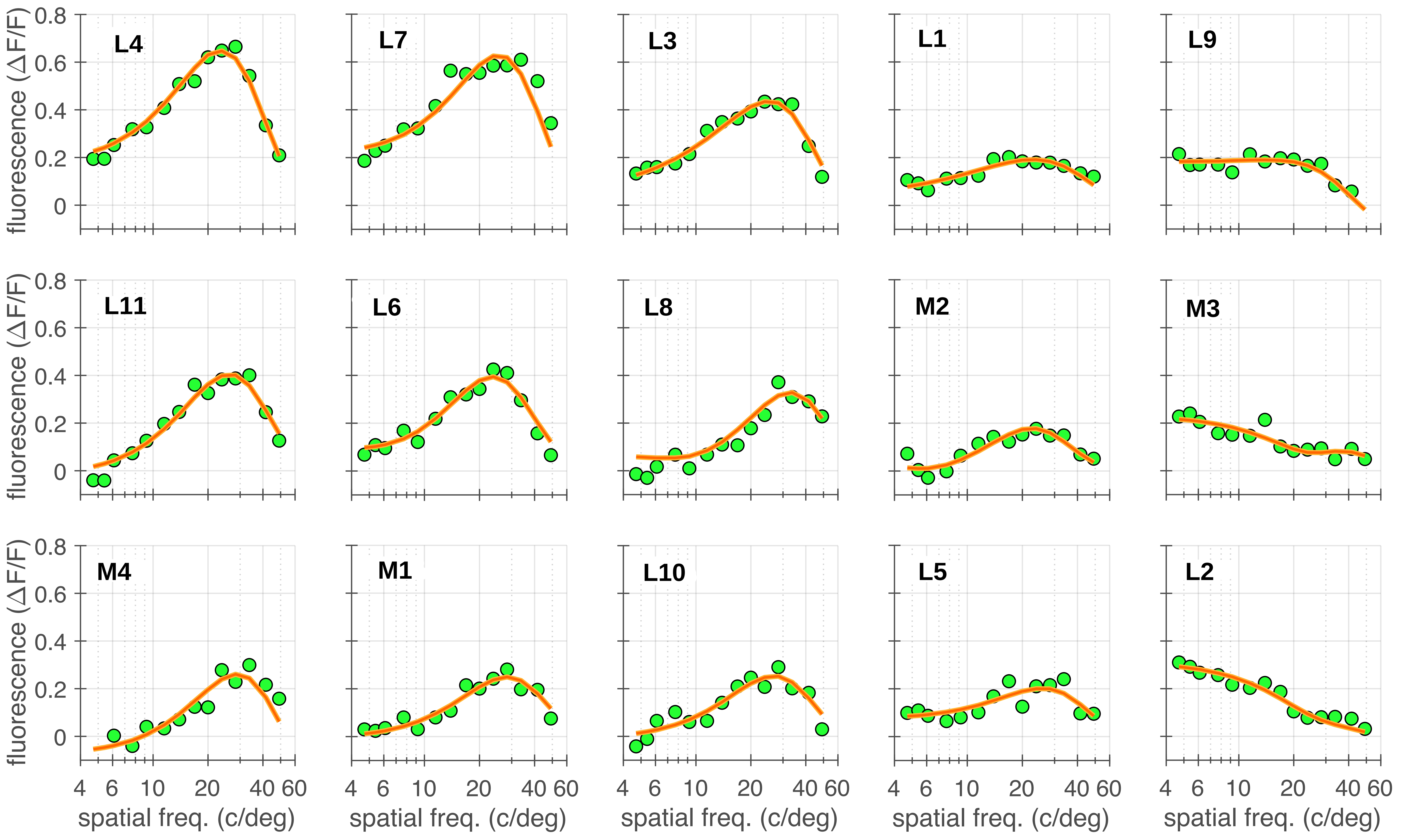


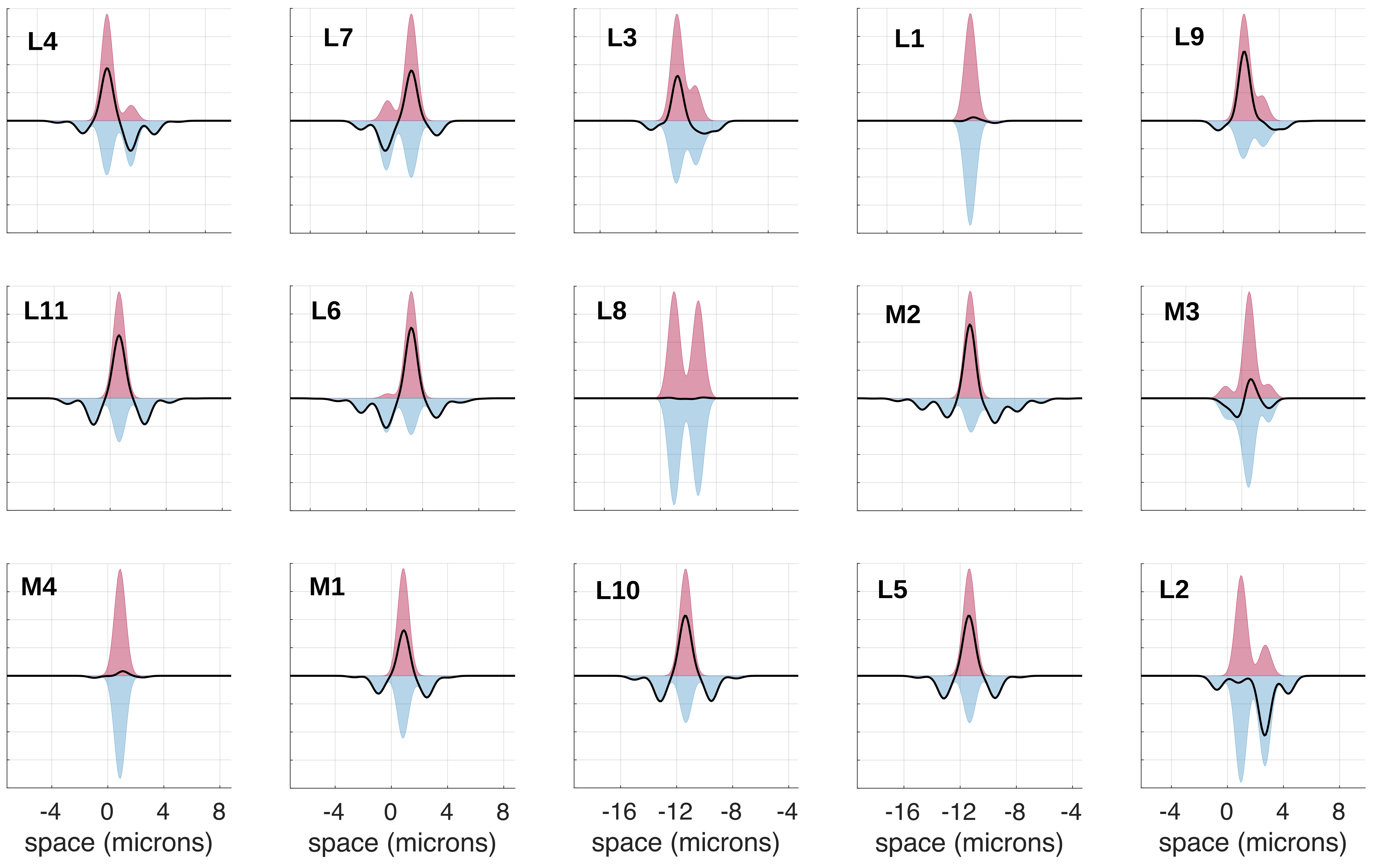
