## Supplement 11 for "*In vivo* chromatic and spatial tuning of foveolar retinal ganglion cells in *Macaca fascicularis*"

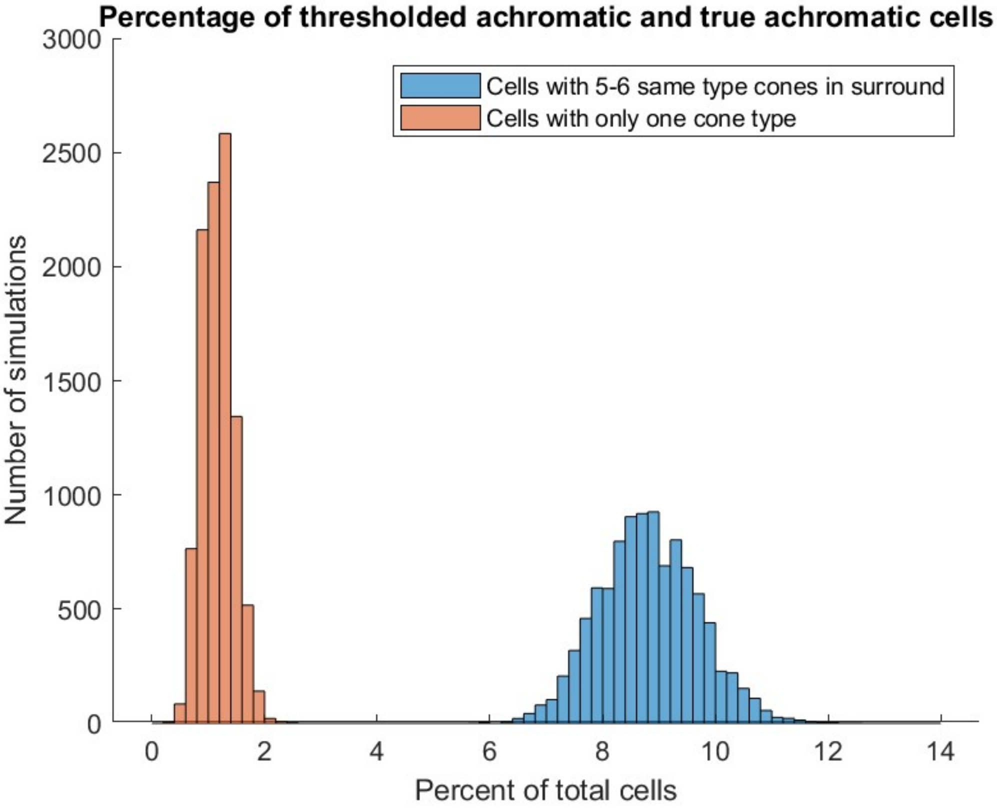


**S11 Fig. Simple center-surround model estimating percentage of achromatic midget RGCs.** A simple model cone mosaic using hexagonal packing was created in MATLAB for the centermost 2400 cones (radius of 25 cones from the foveal center). Each cone was randomly assigned to be an L cone (48%), M cone (48%), or S cone (4%). For this simple model, a midget RGC was assumed to connect to one cone at its receptive field center and 6 cones at its surround (immediately adjacent to the center cone). For each model midget RGC, the proportion of cones that were the same type as the center cone was calculated. RGCs with all 6 surround cones of the same type as the center were classified as 'true achromatic' (orange histogram), and RGCs with at least 5 surround cones of the same type as the center were classified as 'thresholded achromatic' (blue histogram). The entire model simulation was then repeated 10,000 times, with a different randomly assigned cone mosaic each time. The histograms above show the number of simulations for which each percentage of midget RGCs that were either true achromatic or thresholded achromatic occurred. Based on this simple model, truly achromatic midget RGCs might make up approximately 1.5% of total midget RGCs at the foveal center, while thresholded achromatic midget RGCs might make up approximately 9% of total midget RGCs at the foveal center.
